# Determining the rate-limiting processes for cell division in *Escherichia coli*

**DOI:** 10.1101/2024.03.01.582948

**Authors:** Jaana Männik, Prathitha Kar, Chathuddasie Amarasinghe, Ariel Amir, Jaan Männik

## Abstract

A critical cell cycle checkpoint for most bacteria is the onset of constriction when the septal peptidoglycan synthesis starts. According to the current understanding, the arrival of FtsN to midcell triggers this checkpoint in *Escherichia coli*. Recent structural and *in vitro* data suggests that recruitment of FtsN to the Z-ring leads to a conformational switch in actin-like FtsA, which links FtsZ protofilaments to the cell membrane and acts as a hub for the late divisome proteins. Here, we investigate this putative pathway using *in vivo* measurements and stochastic cell cycle modeling. Quantitatively upregulating protein concentrations and determining the resulting division timings shows that FtsN and FtsA numbers are not rate-limiting for the division in *E. coli*. However, at higher overexpression levels, they affect divisions: FtsN by accelerating and FtsA by inhibiting them. At the same time, we find that the numbers of FtsZ in the cell are rate-limiting for cell divisions in addition to other processes. Altogether, these findings suggest that instead of FtsN, FtsZ protofilaments drive the conformational switch of FtsA and lead to the onset of constriction. Our data is also suggestive that FtsA minirings are not present at any significant numbers in wild-type cells.

## Introduction

In most known bacterial species, cell division starts with the formation of the cytokinetic ring, the Z-ring, at the cell’s middle ^1,2^. In *Escherichia coli*, the Z-ring consists of FtsZ protofilaments anchored to the inner membrane via FtsA and ZipA linker proteins. FtsZ protofilaments in the cytokinetic ring are dynamic, undergoing a treadmilling motion ^3,4^. TEM and super-resolution imaging have shown that the Z-ring consists of a sparse set of loosely associated filaments which are distributed in a narrow band (50 nm) around the midcell ^5,6^. Formation of the Z-ring depends on growth conditions starting mostly at cell birth in fast growth rates but being delayed when cells are grown in a poorer quality medium ^7^. Despite the early presence of the Z-ring, the cells do not initiate the onset of constriction concurrent with the formation of the Z-ring. Instead, there is a delay that lasts about ¼ of the cell cycle in *E. coli* for a range of different growth conditions ^8,9^.

While the Z-ring is a prerequisite for cell division, the onset of constriction acts as an effective cell cycle checkpoint ^10,11^. Despite this significance, the signal that triggers the onset of constriction in *E. coli* and other bacteria has remained undetermined. Prior to the onset of constriction, a number of essential divisome proteins are recruited to the midcell. The order of recruitment of these proteins has been determined to be FtsE-FtsX and FtsK→FtsQ-FtsB-FtsL→FtsW-FtsI→FtsN ^12^. Depletion or inactivation of upstream proteins prevents the recruitment of all downstream components. FtsN is the last essential component to arrive at the divisome. Accordingly, it has been considered the trigger protein for cell division in *E. coli* ^13-21^.

During the onset of constriction, a core divisome complex that includes the peptidoglycan synthesizing unit FtsWI ^22^ separates from the treadmilling FtsZ protofilaments ^23^. It has been proposed this separation and subsequent peptidoglycan synthesis is due to the binding of FtsN to the FtsBQL complex that, in turn, activates FtsWI ^19,24^. Before separation, FtsBQL and FtsWI follow FtsZ protofilaments ^23^, likely by the diffusion-and-capture mechanism ^25^. As an alternative pathway, it has been proposed that the change in the oligomerization state of FtsA triggers the checkpoint ^1,26,27^. FtsA, which shares its fold with actin, forms oligomeric structures *in vitro* ^28^. These structures include 12-mer minirings, curved arcs, and antiparallel double filaments ^20,29,30^. Of these higher-order structures, the experimental evidence *in vivo* exists only for the FtsA antiparallel filaments ^20^. It has been proposed that the onset of constriction is triggered by the transformation of FtsA minirings to antiparallel filaments ^1^ or, alternatively, that mostly monomeric FtsA is driven to antiparallel filament form as a result of FtsN binding ^20^.

While the protein-protein interactions between FtsN, FtsA, and other divisome proteins are needed to trigger the onset of constriction, it has remained unclear what causes the switching on of these interactions during the cell cycle. There is no evidence of phosphorylation nor any other post-translational or slow conformational changes of these proteins. As such, the involved protein-protein interactions remain immutable in the cell cycle. On the other hand, *E. coli* and other bacteria have been observed to add approximately a constant length increment from one DNA replication initiation to the next ^31^ and from birth to division, irrespective of their birth length ^32,33^. The latter phenomenon, referred to as the cell size adder, can be explained if cell division is triggered when a number of some protein in the cell reaches a threshold value ^31,34,35^. Most different protein species in the cell appear to increase exponentially during the cell cycle, being synthesized in proportion to cell volume. In principle, a cell could use one of these protein species as a proxy for all of them to determine that its content is approximately doubled before committing to division. A caveat here is that a cell cycle checkpoint is not at the division but at the onset of constriction ^11,36^, and no cell size adder between consecutive constriction events has been reported.

In addition to approximately doubling its protein content, the cell needs to have a mechanism to verify that it has at least two copies of chromosomal DNA. Accordingly, one would expect the onset of division to be tightly coupled to the replication cycle ^31,34,37-39^. However, the tight coordination between replication and division cycles has been disputed ^10,35,40^. Recent experiments and analysis have reconciled these views, indicating that the replication status appears to be only one of the input signals for the onset of constriction, and its effect becomes smaller at faster growth rates ^11,36^. What could then be the driving signal for the onset of constriction at faster growth rates? *Si et al*. proposed that the accumulation of FtsZ to a threshold number could explain the adder correlations in cell length from birth to division in all growth conditions ^35^. However, their work did not quantify the downregulated FtsZ amount. It is expected that a sufficient downregulation of any of the essential divisome proteins and likely also non-essential ones would lead to a delay in the onset of constriction, to longer cells and a decrease in the adder correlations. A more conclusive determination of whether the amount of FtsZ or some other division protein is limiting for the cell division is therefore warranted.

Here, we investigate what is the rate-limiting component or process for the onset of constriction in *E. coli* at fast growth rates using high throughput imaging in microfluidic devices and cell cycle modeling. Our data and modeling show that FtsZ is rate-limiting for the onset of constriction, while the trigger protein FtsN is not. However, in more than two-fold excess concentrations, FtsN starts to promote the onset of constriction. Furthermore, our data also show that FtsA is not rate-limiting nor inhibitory in physiological conditions, but it becomes inhibitory at 50% overexpression level. The latter finding is suggestive that FtsA minirings are not present in physiological conditions.

## Results

Our goal in this work was to develop a quantitative approach to determining if a protein is rate-limiting for cell division. A decrease in concentration/protein numbers of the rate-limiting component should lead to delays in the cell division. However, a sufficiently large down-regulation of an essential divisome component leads to the cessation of the division process even if this component is not rate-limiting in normal growth conditions ^41^. The downregulating concentration of an essential but not rate-limiting protein for cell division, by, say, 20% may produce a delay in the division process, but at 10%, downregulation may still not be limiting. Controlled and quantitative downregulation of this level of accuracy has not been achieved yet. Therefore, instead of downregulating, we rationalized that upregulation of the protein of interest should produce a more robust way of assessing if it is rate-limiting for cell division. An upregulation by any amount should speed up cell division if the component is ratelimiting and have no effect if the component is not rate-limiting. The caveat here is that too high upregulation is known to lead to cytotoxic effects. For example, significant overexpression of FtsA and FtsZ alone (more than 7-fold) leads to cessation of cell division ^42,43^. Upregulation by a small amount and quantification of this amount is thus necessary. Furthermore, attaching a fluorescent fusion for the protein of interest to quantify its concentration/abundance may affect its function so that the altered protein may become rate-limiting due to its impaired function. Having these considerations in mind, we upregulate in our experiments unlabeled protein in wild-type cells (the strain of interest – SOI) and use a strain with a fluorescent proxy on the same chip that expresses a fluorescent protein from the same locus under the same promoter (Reporter strain) (Fig. 1A). To calibrate the signal from the fluorescent Reporter, we also added a third strain (Reference) which expressed the protein of interest fused to the same fluorescent protein as present in the Reporter strain. The Reference strain expresses this fusion protein constitutively from the native locus of the protein of interest. Using real-time signals from Reporter and Reference allows the calibration of upregulated protein amounts in terms of the native protein concentration (for details, see Methods, *Determining Induced Protein Concentrations*). We mixed these three cell populations (SOI, Reporter, and Reference) and grew cells on a microfluidic mother-machine platform (Fig. 1B, C). After growing the cells for about six generations, we upregulated the protein of interest and monitored the change in protein concentration in the cells (Fig. 1D), cell length at division, *Ld*, (Fig. 1E), and the doubling time, *Td*, (Fig. 1F) as a function of time from the upregulation of the protein of interest. We also determined timing, *Tc*, and the cell length, *Lc*, at the onset of constriction (SI Fig. S1).

**Figure 1.**
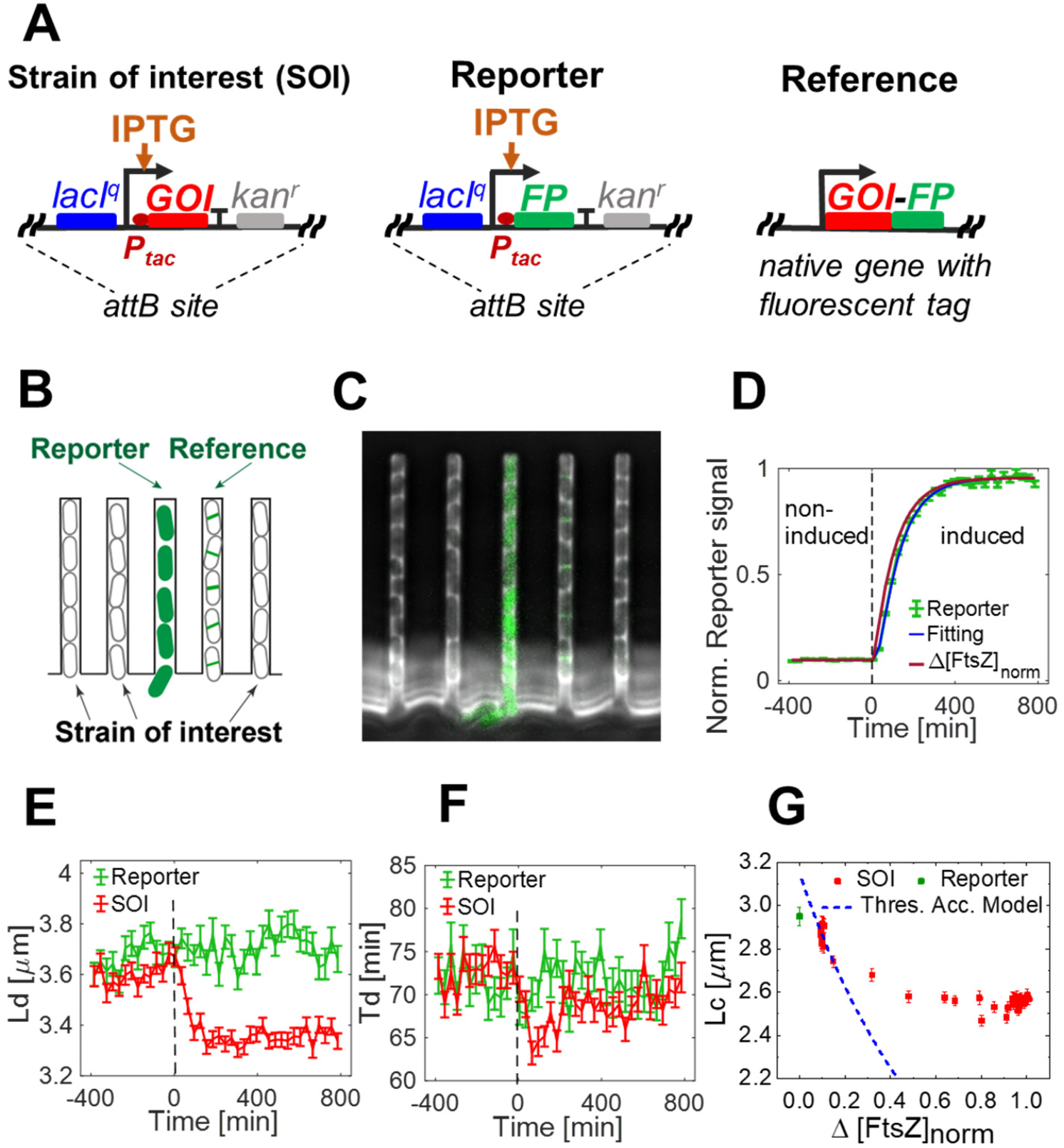
Determining proteins that are rate-limiting for cell division. (A) Experimental design. The Strain of interest (SOI), Reporter, and Reference strains were measured simultaneously to determine the effect of upregulation on cell cycle parameters and to quantify the amount of protein produced. SOI expresses an unlabeled gene of interest (GOI) and the Reporter, a fluorescent protein (FP) from the same promotor. The Reference strain expresses a native protein of interest with the same fluorescent fusion as the Reporter. For a list of all SOI, Reference, and Reporter strains used in experiments see SI Table S4. (B) The strains were mixed and grown on the same microfluidic mother-machine device. (C) A snapshot image of the device after upregulation of the protein of interest, which in this image is FtsZ. (D) Increase of fluorescent reporter signal (Reporter) due to its induction. Addition of IPTG inducer (500 μM) at time zero (indicated by a dashed vertical line). The curve of the Reporter is normalized by the Reference signal. In this case, the overexpression level is 0.96x (96%) in terms of protein concentration in WT cells. The blue line is fitting of the model to the data and the dark red line is the calculated excess normalized concentration of FtsZ, which is corrected for fluorophore maturation (for details, see Methods, *Determining Induced Protein Concentrations*). (E) Change of cell length at division as a function of time for SOI and Reporter. (F) Change of doubling times for the same cells. Time zero corresponds to the addition of inducer IPTG (indicated by a dashed vertical line). (G) Cell length at the onset of constriction versus the excess normalized concentration of FtsZ. The red points show data from SOI, and the green point corresponds to the Reporter strain. The dashed line represents the threshold accumulation model. All error bars correspond to std errors.

The data in Fig. 1E-F and SI Fig. S1 is pertinent to the upregulation of FtsZ from an extra copy from the *λ*-attachment (*attB*) site in moderately fast growth conditions in a glucose-CAS medium. FtsZ upregulation by about 90% leads to a step-like decrease in cell length by about 8% (Fig. 1E) and a transient decrease in doubling time, with the shortest doubling times being 13% shorter than before the protein upregulation (Fig. 1F). The observed speed-up in cell division frequency is qualitatively consistent with the idea that FtsZ is rate-limiting for cell division in *E. coli* at fast growth rates. However, the measurement also shows that at higher overexpression levels FtsZ ceases to be rate-limiting because changes in cell length stop about 100 min after the start of the induction while the FtsZ concentration continues to increase further for about 200 min. The cessation of FtsZ being rate-limiting during the upregulation can be more clearly seen by plotting cell length as a function of the excess normalized FtsZ concentration, *Δ[FtsZ]*_*norm*_, (Fig. 1G). Note that *Δ[FtsZ]*_*norm*_ is based on the normalized reporter signal, which is corrected for fluorophore maturation effects (see Methods). Here, we chose to measure the cell length at the onset of constriction rather than at the division because the former is the cell cycle checkpoint. The data show that as *Δ[FtsZ]*_*norm*_starts to increase, the cell length at constriction decreases linearly in time, but when *Δ[FtsZ]*_*norm*_ exceeds about 70% of the native level, the increase stops. This is approximately the point when FtsZ ceases to be rate-limiting for triggering the onset of constriction for all the cells in the cell population.

To explain the changes in cell length upon FtsZ upregulation quantitatively, we compared the data to the FtsZ threshold accumulation model proposed earlier ^35^ but with a modification that the FtsZ number threshold is reached at the onset of constriction rather than at the division (Fig. 1G). Assuming the FtsZ concentration in this growth condition is constant during the cell cycle yields a simple prediction *Lc =Lc*^*WT*^/(1 *+ Δ[FtsZ]*_*norm*_) as indicated by a dashed line in Fig. 1G. Here *Lc*^*WT*^is the cell length in WT cells without extra FtsZ. The prediction differs in two aspects from the experimental data: 1) the predicted line does not show a plateau at higher *Δ[FtsZ]*_*norm*_ values and 2) the slope of the curve at a small *Δ[FtsZ]*_*norm*_ is much higher than in the data.

### The concurrent process model approximately explains the data

The plateau region in cell length data indicates that in addition to FtsZ accumulation, there could be an additional process that controls the onset of the constriction. This idea is captured by the concurrent processes model ^36,44,45^, where one of the limiting processes stems from FtsZ numbers reaching a threshold value in the cell, and the other limiting process originates from the DNA replication cycle. To compare the model to the experimental data, we used cell cycle parameters determined from the experiments and adjusted the threshold level for the FtsZ numbers and the time duration when the replication-related processes block the onset constriction using the measurement shown in Fig. 1D-G (for details see Methods, *Modeling*). After these adjustments, the model quantitatively explained the increase in fluorescent protein concentration from the Reporter (Fig. 2A), step-like decrease in cell length (Fig. 2B, SI Fig. S2A), and the transient decrease in timings for the onset of constriction (Fig. 2C) and cell division (SI Fig. S2B). Note that to reproduce the change in steady state timings for the constriction (*Tc*) we needed to account for the increased constriction period (*ΔTs = Td - Tc*) that resulted from FtsZ upregulation (SI Fig. S3).

**Figure 2.**
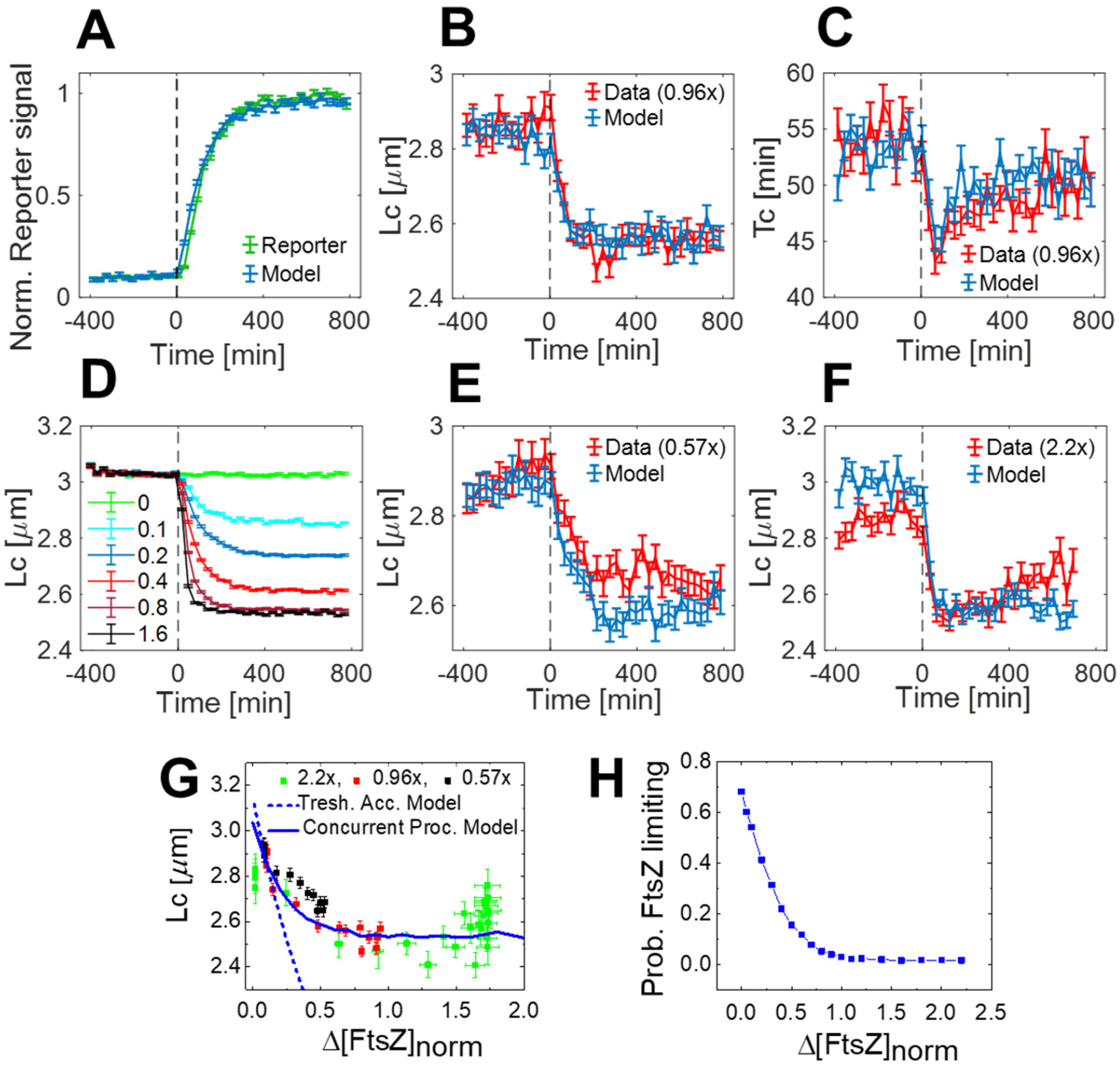
Comparing the experiment and the concurrent processes model. (A-C) The normalized reporter signal, the cell length, and the timing of the onset of constriction from the experiment (red) and model (blue). The experimental curves correspond to the measurement in Fig. 1. (D) Model predictions for cell length at the onset of constriction for different levels of upregulation of FtsZ. The legend shows the final steady-state value of *Δ[FtsZ]*_*norm*_ for each curve. (E-F) Comparing model predictions for *Lc* to measurements where the final steady-state value is *Δ[FtsZ]*_*norm*_ *=* 0.57 and *Δ[FtsZ]*_*norm*_ *=* 2.20, respectively. For comparisons of other measured quantities, see SI Figs. S4 and S5. (G) *Lc* vs. *Δ[FtsZ]*_*norm*_ from three FtsZ upregulation measurements combined. The blue solid line is the prediction from the concurrent process model, and the dashed line is from the threshold accumulation model. (H) The probability that FtsZ numbers limit the onset constriction in a cell population according to the concurrent processes model.

Qualitatively, the cell length and transient decrease in timings for the onset of constriction and division can be understood in the model because, at the higher synthesis rate of FtsZ, the cells reach the number threshold earlier in the cell cycle when their length is smaller. However, as the cells get smaller, the synthesis rate of FtsZ, which is proportional to cell volume/length, decreases. The timings *Tc* and *Td*, therefore, shift back to their original values after the completion of one whole cell cycle that follows the upregulation of FtsZ (without accounting for increased *ΔTs*). The unchanged steady state timings *Tc* and *Td* after upregulation of FtsZ from their pre-induction values can also be understood from the constancy of cell growth rates before and after upregulation. However, as mentioned above, we adjusted the basic model for the increased constriction period (*Td - Tc*), which accompanies FtsZ upregulation. The increased constriction period shortens both *Tc* and *Td* because more FtsZ accumulates during the longer constriction period in the previous cell cycle.

To test the model further, we carried out additional experiments where FtsZ concentration at the final steady state differed from the one shown in Fig. 2A-C. The model predicts that because of the increase in the final FtsZ steady-state concentration, the change in cell length at constriction approaches a minimal limiting value, and the duration of cell length change decreases (Fig. 2D). The minimal limiting length is determined by replication-related processes alone. Without any further parameter adjustments, the model qualitatively predicted the time-dependences of all measured quantities in both overexpression conditions (Fig. 2E-F, SI Fig. S4, S5). At the quantitative level, however, the experimentally observed effect was smaller than the model prediction for the cell lengths at the onset of constriction in the low overexpression conditions (0.57x) (Fig. 2E). The same was true also for the higher (2.2x) overexpression measurement (Fig. 2F). However, in this measurement, the effect appeared as incorrectly predicted *Lc* value before upregulation. This is because, before the upregulation of FtsZ, there was a much smaller leaky expression of FtsZ in high (2.2x) overexpression measurement (*Δ[FtsZ]*_*norm*_ *=*0.02) compared to the two lower FtsZ overexpression measurements (*Δ[FtsZ]*_*norm*_ *=*0.10). The model was thus unable to quantitatively predict the effects arising from this leaky expression. The failure of the model to correctly predict cell length due to leaky expression of FtsZ was also evident when we compared the cell lengths for the Reporter and SOI strains (SI Fig. S6). The behaviors in all overexpression levels can be summarized in the FtsZ titration plot (Fig. 2G). The model can be seen to fit the data well except for the *Δ[FtsZ]*_*norm*_ ≲ 0.1 and for *Δ[FtsZ]*_*norm*_ ≳ 1.3. The deviation of the model for *Δ[FtsZ]*_*norm*_ ≳ 1.3 (130% overexpression) can be explained by the cytotoxic/inhibitory effects of FtsZ overexpression, as reported earlier (42).

While the deviation of the model from the experiments for *Δ[FtsZ]*_*norm*_ ≳ 1.3 was expected, the discrepancy at *Δ[FtsZ]*_*norm*_ ≲ 0.1 presents a more significant challenge for the model. A key characteristic of the concurrent processes model is a large cell length change for variations in FtsZ concentration close to *Δ[FtsZ]*_*norm*_ *=* 0 and decrease of this response as *Δ[FtsZ]*_*norm*_ increases (Fig. 2G). Such behavior is independent of model parameters. Failure of this prediction suggests that a number threshold postulated by the model may not accurately capture cellular response near the WT concentration of FtsZ. Instead of a simple number-sensing mechanism, there could be a more complex response function involved in the decision-making process that triggers the onset of constriction. At the same time, the model fits the data well at high overexpression conditions, suggesting that at a coarser level, the accumulation to threshold number provides a good, effective model to explain how the onset of constriction is triggered.

### Quantifying FtsZ limitation

A comparison of data and model shows that the FtsZ amount in the cell is one of the rate-limiting factors for the onset of constriction. The question then arises: how limiting is FtsZ? As mentioned above, the concentration-dependent decrease in cell length plateaued at about 0.7x (70%) overexpression levels of FtsZ (Fig. 2G) indicating that at this overexpression level, FtsZ is effectively not limiting the onset of constriction for any cell in a population. Alternatively, the concurrent processes model allows us to estimate the probability that in a given division FtsZ numbers are limiting the onset of constriction. This probability as a function of *Δ[FtsZ]*_*norm*_ is shown in Fig. 2H. The probability approaches zero at *Δ[FtsZ]*_*norm*_» 0.7 as expected from the previous argument. More importantly, the model predicts that FtsZ is limiting in 68% divisions for the WT cells at the native level of FtsZ (i.e. *Δ[FtsZ]*_*norm*_ *=* 0). It is worth emphasizing that this number is as valid as is the model. Since the model shows a smaller change in cell lengths than the experiment at low overexpression levels of FtsZ, 68% probability is likely an overestimate. Nevertheless, we can draw a rough estimate that for at least half of the cell divisions in WT cells in this moderately fast growth condition, FtsZ numbers in the cell rate limit the onset of constriction.

### FtsN is not rate-limiting for the onset of constriction

Our modeling considered replication the second limiting process, but similar results can be obtained if, instead of replication, another division-related protein limits the onset of constriction (SI Fig. S7). Several recent works have argued that FtsN acts as a trigger for the onset of constriction ^13-15,17-20,46^. Therefore, we also carried out the upregulation measurements for FtsN. In the first set of measurements, we upregulated FtsN concentration by about 70% (Fig. 3A), which is similar to the FtsZ overexpression level in Fig. 1. Unlike for FtsZ, there was no change in *Ld, Lc, Td* or *Tc* upon overexpression of FtsN (Fig. 3B-C, Fig. S8A-B). This data thus rules out that FtsN is a rate-limiting for cell division in these conditions. However, when we repeated these measurements at a much higher upregulation level (6.5x, Fig. 3D), then a clear decrease in cell length appeared (Fig. 3E, SI Fig. S8C-D). Combining these measurements in the FtsN titration curve showed that cell length is insensitive to upregulations for *Δ[FtsN]*_*norm*_ ≲ 2.0 but above this value it starts to decrease (Fig. 3E-F). These measurements thus show that FtsN is not rate-limiting to cell division at its native levels. At higher concentrations, it can accelerate the onset of constriction as observed earlier ^47^.

**Figure 3.**
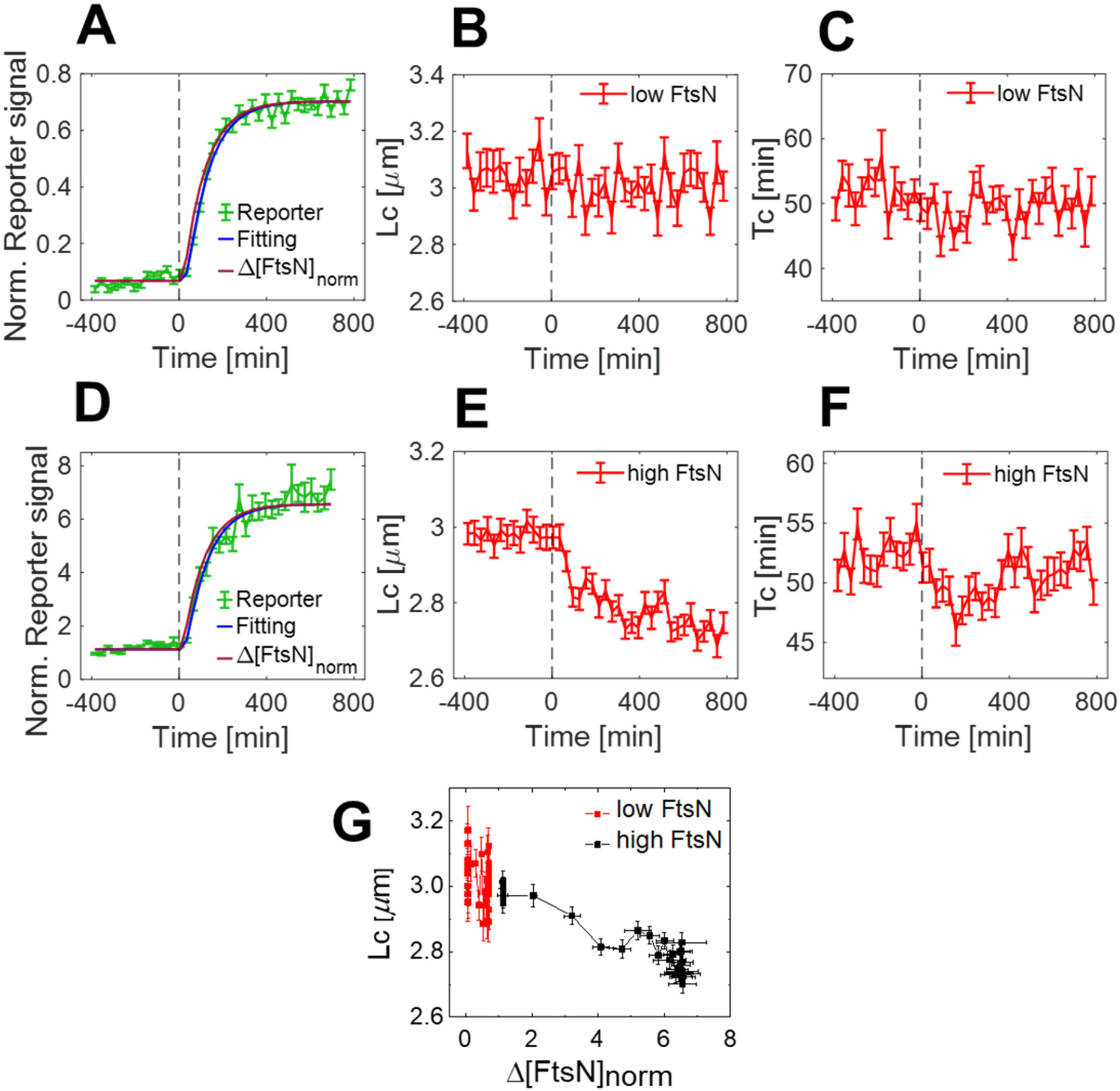
Upregulation measurements of FtsN. (A) Increase of fluorescent reporter signal (Reporter) due to its induction at a low level of upregulation (0.71x). The curve is normalized by the Reference signal. The blue line fits the model to the data, and the dark red line is the calculated total upregulated protein concentration. (B-C) The cell length at the onset of constriction and the timing for the onset of constriction as a function of time, respectively. The final steady-state value *Δ[FtsN]*_*norm*_ *=* 0.71 (comparable to FtsZ upregulation measurement in Fig.1). Time zero corresponds to the addition of inducer IPTG (indicated by a dashed vertical line). (D-F) The same measurements as in the top row but for an induction at a high level of upregulation (6.56x). (G) *Lc* vs. *Δ[FtsN]*_*norm*_ from these two FtsN upregulation measurements combined. All error bars correspond to std errors.

### A low level of FtsA overexpression is not inhibitory

FtsA is the second most conserved bacterial cell division protein besides FtsZ ^48^. At several-fold upregulation, FtsA is cytotoxic, preventing the onset of constriction ^43,49^. However, it is possible that at native levels, it is rate-limiting for cell division like FtsZ because the two proteins are expressed from the adjacent genes in the same operon. To test this hypothesis, we carried out overexpression measurements also for FtsA (Fig. 4A). Contrarily to our hypothesis, FtsA in low overexpression conditions (the final *Δ[FtsA]*_*norm*_ *=* 0.60) did not lead to a change in cell length (Fig. 4B, SI Fig. S9A-B), suggesting that FtsA is not rate-limiting for cell division in normal growth conditions. However, at higher overexpression levels of FtsA (the final *Δ[FtsA]*_*norm*_ *=* 1.5) led to an increase in cell length (Fig. 4C; SI Fig. S9C-D). This increase only occurred after a distinct delay, confirming that at the low level of overexpression, FtsA is not inhibitory. The increase in cell length at higher overexpression levels is consistent with earlier reports ^9,49^. Converting the time-dependent measurements to the FtsA titration curve showed that FtsA inhibitory effects set in at about 60-70% overexpression levels (Fig. 4D). It is worth mentioning that 60-70% overexpression of [FtsA] beyond its mean value is likely beyond the natural variation in a cell population. So, in a natural population, FtsA is not expected to be inhibitory for the vast majority or even in any cells in the population.

**Figure 4.**
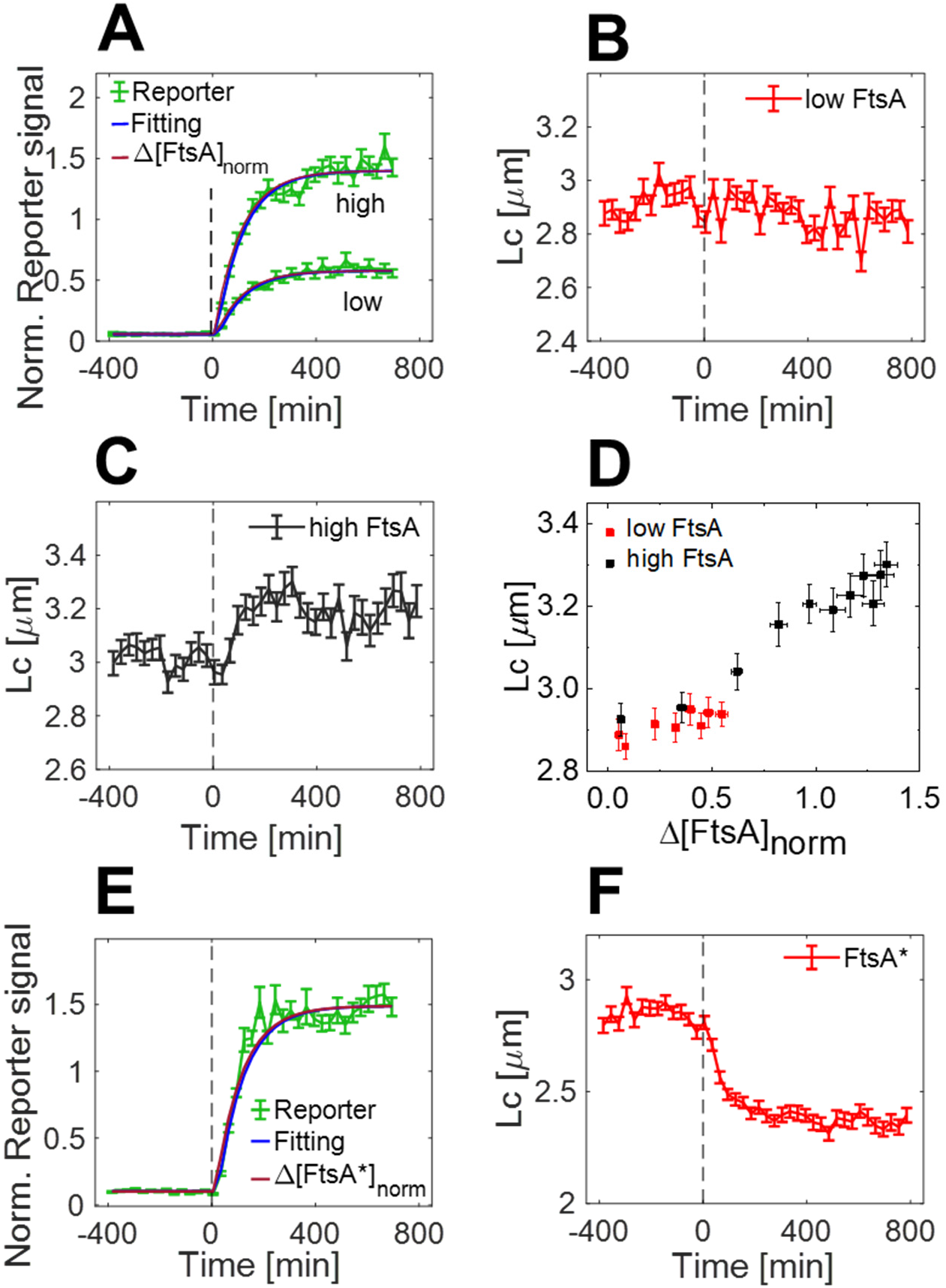
Effects of overexpression of FtsA and FtsA*(R286W). (A) Increase of fluorescent reporter signal (Reporter) as a result of its induction at a low level of upregulation (60% excess, low) and a high level of upregulation (150% excess, high). The blue line is fitting of the model to the data and the dark red line is the calculated total upregulated protein concentration. (B,C) Cell length at the onset of constriction vs time for the final *Δ[FtsA]*_*norm*_= 0.6 and 1.5, respectively. (D) The average cell length at the onset of constriction versus *Δ[FtsA]*_*norm*_ from the two measurements combined. (E) Increase of fluorescent reporter signal (Reporter) as a result of its induction. The blue line is fitting of the model to the data and the dark red line is the calculated total upregulated protein concentration. F) The cell length at the onset of constriction for the final *Δ[FtsA*^∗^]_*norm*_ *=* 1.5. All error bars are std errors.

Unlike FtsA, overexpression of its hypermorphic mutant FtsA* (R286W) is not known to cause cytotoxic effects and leads to short cell phenotype ^50^. To quantify its effects, we also upregulated FtsA* in the WT FtsA background (about 1.5x; Fig 4E). Upregulation led to decreased cell length (Fig. 4F, SI Fig S10A-D) and a transient decrease in the timing for the onset of constriction (SI Fig S10B). The effect appeared very similar to the ones from FtsZ upregulation measurements. We overlayed the two measurements (SI Fig. S10 E-H), finding the effects of FtsZ and FtsA* upregulation almost indistinguishable from each other. Note the Ld vs. Time and Td vs. Time curves are insensitive to the amounts of protein upregulation in the range of concentrations (cf. Fig. 2D). Since FtsZ and FtsA* have the same downstream effect, they may cause the same change in the divisome that is needed to trigger the onset of constriction.

## Discussion

Key proteins in the cell division in *E. coli* are the earliest-arriving components FtsZ and FtsA, and the last arriving essential protein FtsN. Here, we investigated if any of them are rate-limiting for cell division by modestly upregulating their concentrations. Although the upregulation of these proteins has been carried out in the past, the increase of protein concentrations in these measurements far exceeded the physiologically relevant range and has not been precisely quantified. The data presented here shows that FtsZ is rate-limiting for cell division at moderately fast growth conditions, while FtsN and FtsA are not.

While the rate-limiting role of FtsZ is expected, the finding that FtsN is not rate-limiting refutes the idea that it is the trigger protein. This notion does not mean that FtsN is not needed/essential for the onset of constriction. Based on the terminology proposed earlier ^41^, FtsN is a secondary rather than a primary cell cycle regulator. Clearly, depletion of FtsN leads to cessation of cell division. However, it is present in the cell in excess concentration and ready to be incorporated into the divisome complex early on in an FtsA* background as was demonstrated earlier ^9^. The results from this earlier work, thus, showed that FtsN is not sequestered away by some yet-to-be-identified protein interactions in earlier stages of the cell cycle or undergoing some post-translational modifications before the onset of constriction. Instead, there is some other change in the upstream complex that needs to occur before FtsN can bind to the complex. Rather than being a trigger to the onset of constriction, the binding of FtsN to activated complex appears to be a consequence of some other event occurring before FtsN binds to FtsA.

What could this event be? Our data indicates that increased FtsZ concentration drives this transition. We used a previously published model ^51^ to understand what effect an increase in FtsZ concentration has on FtsZ protofilaments. Based on modeling, the excess FtsZ leads to an increased number of FtsZ protofilaments (SI Fig. S11). The increase in filament number is approximately proportional to the increase in FtsZ concentration. At the same time, the increase in concentration changes the filament length only weakly. More filaments of approximately the same length can be expected to lead to elevated filament bundling. Based on these arguments, we can infer that FtsZ protofilament bundling is needed for the onset of constriction.

There are several possibilities of what effect FtsZ protofilament bundling may have. A high local concentration of FtsZ protofilaments due to bundling may cause a small inward bending of the plasma membrane, which may be needed to initiate septal peptidoglycan synthesis. The bending would be mediated by FtsA and ZipA. FtsA might have a dominant effect over ZipA because of its shorter disordered linker, which would bear a majority of the load from bending intrinsically curved FtsZ protofilaments ^52^. It remains, however, unclear how the excess of FtsA could hinder and the excess of FtsA* mediate the force transduction. Furthermore, the magnitude of the bending may not be significant *in vivo*, where the turgor pressure could oppose the membrane deformations.

Alternatively, FtsZ protofilament bundling can enforce a conformational change in FtsA, which in turn activates septal peptidoglycan synthesis. Recent data indicate that for the latter to occur, FtsA antiparallel filaments are needed ^20^. The same work found that the transition to antiparallel filaments can be induced by adding high concentrations of FtsN N-terminal cytoplasmic tail to the reaction mixture. Our data indicate that FtsN is not rate-limiting *in vivo*, while FtsZ is. Accordingly, we propose that the transition to FtsA antiparallel conformation is driven by FtsZ but not by FtsN.

*In vivo* measurements indicate that FtsA antiparallel filaments are not thermodynamically favored compared to arcs and minirings when only FtsA alone is present in the reaction mixture ^20,30^. We postulate that for the formation of a FtsA antiparallel filament, FtsZ protofilaments need to form at least a local doublet to nucleate FtsA antiparallel filaments (Fig. 5A). A straight doublet of FtsZ protofilaments may force monomers and curved FtsA arcs to a straight (antiparallel) filament conformation. In this conformation, 1:1 stoichiometry between FtsZ and FtsZ locally holds where every C-terminal tail of FtsZ is bound to FtsA. Based on the proposed model, these FtsZ C-terminal connections to FtsA drive the FtsA antiparallel filament formation. However, FtsZ protofilament doublets need to be parallel to stay together during treadmilling while FtsA oligomers are antiparallel. This appears not to be a contradiction. We expect the free energy difference of FtsZ binding to FtsA parallel strand compared to the antiparallel strand to be minimal because FtsZ has a long and flexible C-terminal linker. Once FtsA antiparallel filament nucleates, it is further stabilized by binding FtsN to it. The nucleation and antiparallel filament formation can also be driven by FtsN alone without FtsZ doublets and bundles but at much higher concentrations of FtsN than present in cells *in vivo* (Fig. 5B). The proposed mechanism works as if the arrival of FtsN to the divisome triggers the onset of constriction even though the actual mechanism is the conformational switch of FtsA induced by FtsZ protofilament bundling. The proposed mechanism does not contradict recent single-molecule tracking data where some fraction of FtsBQL and FstIW complexes were found to move together with treadmilling FtsZ protofilaments ^21,23^. These complexes can bind to mostly monomeric FtsA and follow treadmilling FtsZ protofilaments by diffusion and capture ^25^.

**Figure 5.**
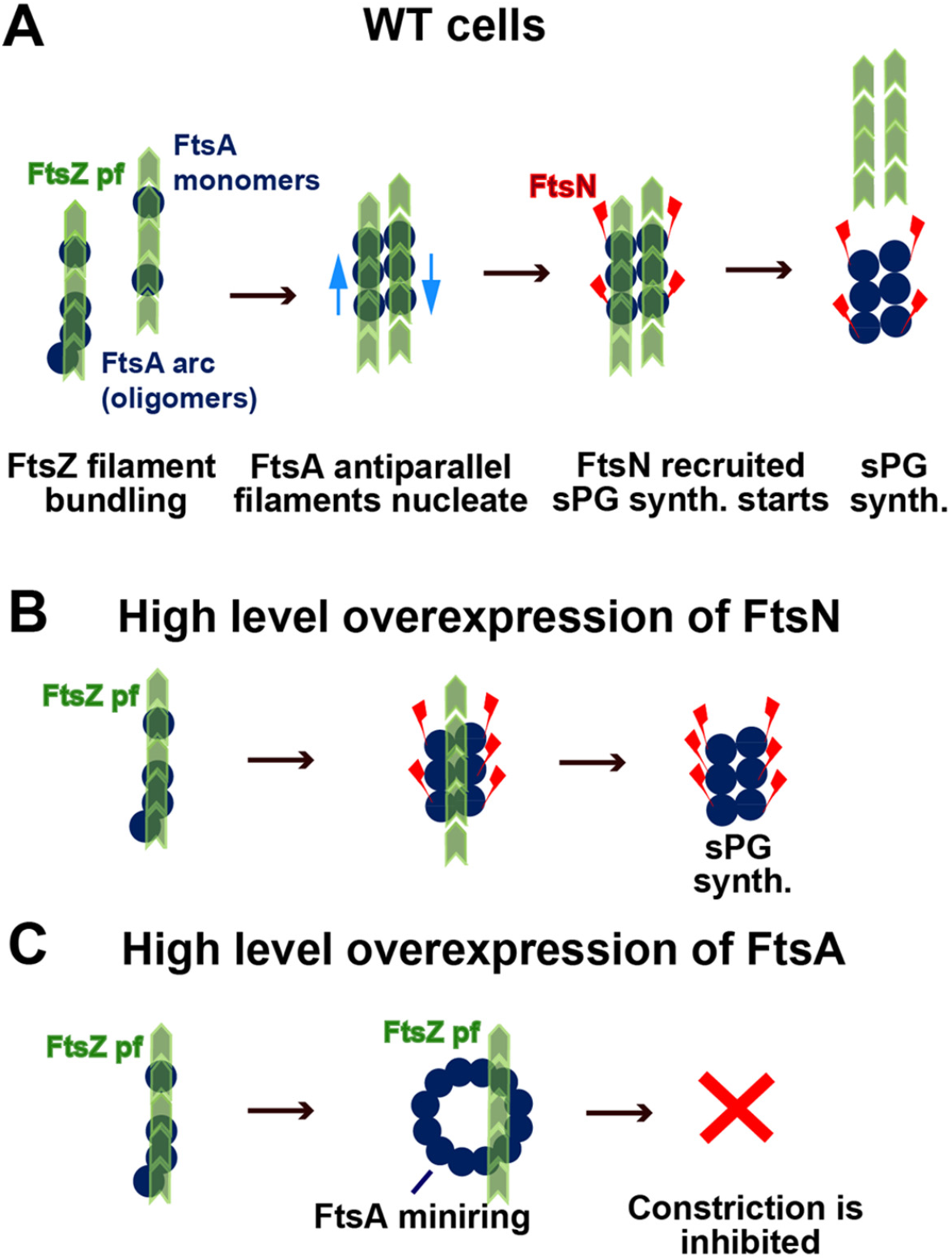
A proposed mechanism for the onset of constriction. (A) In WT cells, an FtsZ protofilament doublet or a larger bundle is needed to nucleate the FtsA antiparallel filament. The nucleated filament is stabilized by FtsN. (B) Unnaturally high concentrations of FtsN can lead to FtsA antiparallel filament formation without FtsZ protofilament doublet. (C) Unnaturally high concentrations of FtsA can lead to FtsA minirings, which prevent FtsZ protofilament bundling and inhibit constriction formation.

Alternatively, one could consider a mechanism where the onset of constriction is triggered by the transition of FtsA minirings to its antiparallel filament conformation (SI Fig. S12) ^1^. The only difference between the two mechanisms is the initial polymerization state and confirmation of FtsA (monomers and FtsA arcs vs minirings). FtsA minirings can be expected to be inhibitory for the onset of constriction because of the occlusion of the FtsW binding interface in this conformation of FtsA ^53^. If FtsA minirings were already present in cells in the native expression level of FtsA, then we would expect a continuous increase in cell length as FtsA concentration increases and more and more FtsA minirings form that are not competent for recruiting downstream components. Instead, our data show an inhibitory effect arising from FtsA overexpression only at 60% overexpressed conditions (Fig. 4D). We interpret the onset of inhibitory effects as a threshold for FtsA miniring formation. Consistent with this idea, *in vitro* measurements have also shown a well-defined threshold for the FtsA miniring formation as the concentration of FtsA increases ^27,30^.

FtsA* is defective in forming minirings ^30^, and it was observed to transform more readily to antiparallel filament confirmation *in vivo* in the presence of FtsN ^20^. Expression of FtsA* could accelerate the onset of constriction, as observed in our measurements (Fig. 4F), because it prevents short FtsA arcs from forming, and this facilitates the nucleation of FtsA antiparallel filaments. So, its effect is similar to the one we observe in FtsZ upregulated conditions (SI Fig. S10 E-H). Within this picture, both FtsA* and FtsZ overexpression facilitate the formation of FtsA antiparallel filaments. FtsA* achieves this by shortening FtsA oligomers and thereby lowering the activation barrier for nucleation of FtsA antiparallel filaments.

### Future directions and outlook

Altogether, our data is consistent with the model where an increase of FtsZ numbers in the cell cycle is one of the driving mechanisms for the onset of constriction at faster growth rates. The possible molecular pathway, which is consistent with our and other published data, involves the formation of FtsZ bundles (doublets), which drive FtsA from a monomer and short arc form to an antiparallel filament form. FtsA antiparallel filament form is competent in the recruitment of FtsN, which activates the core divisome complex and starts the onset of constriction.

Although the existing data is consistent with the proposed model (Fig. 5A), alternative explanations cannot be ruled out. To further validate the model in Fig. 5A, the existence of FtsZ protofilament doublets and bundles needs to be verified *in vivo* setting. Also, *in vitro* measurements assessing the role of FtsZ protofilaments on the assembly of FtsA antiparallel filaments would be valuable. Finally, the question also remains on how the core divisome complex, consisting of FtsIW, FtsBQL, and FtsK, is exactly linked to FtsA antiparallel filaments.

## Acknowledgments

The authors thank Harold Erickson for valuable discussions, Scott Retterer for help in microfluidic chip making, and Joe Lutkenhaus and Sebastien Pichoff for plasmids. A part of this research was conducted at the Center for Nanophase Materials Sciences, which is sponsored at Oak Ridge National Laboratory by the Scientific User Facilities Division, Office of Basic Energy Sciences, U.S. Department of Energy. This work was supported by the National Institutes of Health award GM127413 (JM) and NSF MCB2313719 (JM). AA and PK acknowledge support from NSF CAREER 1752024. AA thanks for the generous support of the Clore Center for Biological Physics and ERC-CoG 2023 101125981.

## Author Contributions

JaanM and JaanaM designed the experiments, JaanaM contributed all the bacterial strains and plasmids and carried out the experiments, CA and JaanM designed, and CA fabricated the microfluidic devices, AA and PK developed the model and carried out modeling, JaanM wrote the manuscript. All authors commented on, edited, and approved the manuscript.

## Methods

### Bacterial strains

All *E. coli* strains used in the reported experiments are derivatives of K12 BW27783 obtained from the Yale Coli Genetic Stock Center (CGSC#: 12119). Information on all strains and plasmids are listed in Tables S1 and S2, respectively. Oligonucleotide information is given in Table S3.

For FtsZ and FtsN overexpression measurements, we constructed three strains: 1) the strain of interest (SOI) that overexpressed unlabeled gene of interest in addition to the native one, 2) the reporter strain that expresses fluorescent fusion protein instead of the gene of interest from the same genomic locus, 3) the reference strain that expresses fluorescent fusion to the gene of interest from its endogenous locus.

### Strains for FtsZ overexpression measurements

Firstly, a plasmid pEXT22-FtsZ (pJM142) with unlabeled FtsZ under the control of an isopropyl-β-D-thiogalactoside (IPTG) inducible *P*_*tac*_ promoter was constructed. Briefly, FtsZ was amplified from plasmid JW0093 ^3^ using primers PR200 and PR201. The Ribosome Binding Sequence (RBS, TAGAGAAAGAGGAGAAATACTAG) was introduced in part of the forward primer in front of the *ftsZ* sequence. The resulting insert (*xbaI_RBS_ftsZ_hindIII*) was cloned into vector pEXT22 ^54^ using restriction enzymes *XbaI* and *HindIII*. The same approach was also used for the Reporter plasmid expressing *mNeonGreen* instead of *ftsZ*. In this case, *mNeonGreen* was amplified from plasmid pJM21 (Männik lab) using primers PR202 and PR203. The resulting insert (*xbaI_RBS_mNeonGreen_hindIII*) was ligated into pEXT22 using restriction enzymes *XbaI* and *HindIII* and transformed into DH5α competent cells (Thermo Scientific). After verification by restriction analysis and DNA sequencing, the plasmids were transformed into the BW27783 strain, yielding strains JM237 (FtsZ expressing) and JM228 (mNeonGreen Reporter).

To obtain low-level FtsZ overexpression SOI (JM252) and corresponding Reporter mNeonGreen expressing strain (JM250), the DNA cassette of the *lacIq-p*_*Tac*_*-ftsZ-aph* from pEXT22-FtsZ was introduced into the λ-attachment (*attB*) site by *λ*-Red engineering ^55^. For chromosome insertion, the *lacIq-p*_*Tac*_*-ftsZ-aph* cassette was amplified from plasmid pEXT22-FtsZ using primers (PR206/ PR207), each composed of a 42 (44)-bp sequence at the 5’ end homologous to the *attB* region. The resulting PCR product was treated with *DpnI* (NEB), gel purified using GeneJET Gel Extraction and DNA Cleanup Micro Kit (Thermo Scientific), and then electroporated into a JM217 strain containing pSIM5 ^55^ that encodes λ-Red proteins. Kanamycin-resistant colonies were verified by using colony PCR with primers PR210 and PR211 (or PR212), followed by direct sequencing. The *lacIq-p*_*Tac*_*-ftsZ-aph* allele was then transduced from JM246 into BW27783 by P1 transduction to generate strain JM252. In parallel, the same procedures were conducted to make the Reporter strain expressing mNeonGreen from the *attB* locus. Finally, the *lacIq-p*_*Tac*_*-mNeonGreen-aph* allele was transduced from JM248 into BW27783 to generate strain JM250. The reference strain (JM147) expresses endogenous FtsZ-mNeonGreen^SW^. The same reference strain was also used in the above-described experiments with pEXT22-FtsZ.

### Strains for FtsN overexpression measurements

For FtsN over-expression strains (JM200, JM230), a plasmid expressing Ypet-FtsN (JM200) and unlabeled FtsN (JM230) under the control of an IPTG inducible weakened *P*_*trc*_ promoter was constructed. *Ypet-ftsN* was amplified from the gDNA of the STK13 strain ^11^ using primers PR204 and PR205. The resulting insert (*ecoRI_RBS_ypet-ftsN_hindIII*) was cloned into vector pDSW210 (JM149) using restriction enzymes *EcoRI* and *HindIII*. For unlabeled FtsN expression, *ftsN* was amplified from plasmid pDSW210-Ypet-FtsN using primers PR213 and PR205. The resulting insert (*xmaI_ftsN_hindIII*) was cloned into pDSW210-Ypet-FtsN (JM230) using restriction enzymes *XmaI* and *HindIII*. Also, the reporter strain expressing plasmid pDSW210-Ypet was constructed in a similar way. In this case, *ypet* was amplified from pDSW210-Ypet-FtsN using primers PR204 and PR214. The resulting insert (*ecoRI_RBS_ypet_hindIII*) was cloned into pDSW210-Ypet-FtsN (JM200) using restriction enzymes *EcoRI* and *HindIII*. To be able to compare expression levels of labeled and unlabeled FtsN strains, the same sequence, including RBS, was retained in front of the *ypet* sequence in pDSW210-Ypet (JM241), in front of *ftsN* expressed from pDSW210-FtsN (JM230), in front of *ypet-ftsN* expressed from pDSW210-Ypet-FtsN (JM200) as it is in upstream of native ypet-FtsN in strain STK13.

For low-level overexpression of FtsN and Ypet-FtsN, the DNA cassette sequences from respective plasmid constructs were inserted into the *attB* site by *λ*-Red engineering as described above for the *ftsZ* gene. Briefly, the *lacI*^*q*^*-p*_*Trc*_*-ftsN-bla* and *lacI*^*q*^*-p*_*Trc*_*-ypet-ftsN-bla* cassette was amplified from a template plasmid pJM230 or pJM200, respectively, using primers (PR208/ PR209) and electroporated into JM217 strain with pSIM5 plasmid. Ampicillin-resistant colonies were verified by colony PCR with primer pair PR210/PR211, followed by direct sequencing. In the final step, the resulting alleles were transferred to BW27783 genetic background by P1 transduction generating strains JM235 (inducible unlabeled FtsN at the *attB* site) and JM222 (inducible Ypet-FtsN at the *attB* site).

### Strains for FtsA and FtsA* (FtsA^R286W^) overexpression measurements

Plasmids expressing *ftsA* (pDSW210-ftsA, also known as pSEB306+) and *ftsA\** (pDSW210-ftsA^*^, also known as pSEB306*+) under an IPTG inducible promoter were transformed into BW27783 generating strains JM242 and JM243, respectively. The strain JM149 containing pDSW210-GFP expresses GFP under an IPTG inducible promoter and was used as the Reporter. These three plasmids were described previously ^56^. These upregulation measurements lacked reference strain because of an absence of genomically integrated fully functional FtsA. Instead, Western blotting was used to determine the final *Δ[FtsA]*_*norm*_ for 100 μM IPTG induction ^9,56^ for both FtsA (1.39x) and FtsA* (1.43x). The corresponding calibration factor was also used to quantify the expression level of FtsA in low overexpression conditions (0.43x) where 30 μM IPTG induction was used.

For *E. coli* strain engineering, the cells were grown in lysogeny broth (LB) and appropriate selective antibiotics.

### Growth media and growth conditions

For time-lapse imaging in microfluidic devices, the cells were cultured in M9 minimal media supplemented with 2 mM magnesium sulfate, 0.5% glucose as the carbon source, 0.5% casamino acids (CAS), and 1x trace metals mixture (Tre) at 28°C. When appropriate, the medium was supplemented with 100 μg/ml ampicillin (Amp) or 40 μg/ml kanamycin (Kan). Amp and Kan concentrations were reduced to 20 μg/ml when cells carried *bla* or *aph* integrated into the chromosome. For induction, IPTG (30 μM, 100 μM, 500 μM, or 1 mM) was included in the media.

### Cell preparation and culture in microfluidic devices

The strains were streaked on M9 minimal media agar plates supplemented with magnesium sulfate, glucose, CAS, Tre, and appropriate selective antibiotics. For the microscopy experiment, a colony from a fresh plate was inoculated into 3 ml of M9 minimal salt media supplemented with carbon source and additives described above. 40 μg/ml of Kan was added to grow strains JM147, JM228, and JM237. 100 μg/ml of Amp to grow strains JM149, JM230, JM241, JM242 and JM243. The cells were grown to an OD_600_ of ∼0.1 in a liquid medium and then concentrated ∼100x by centrifugation in the presence of 0.075 μg/ml of BSA (Bovine Serum Albumin; Millipore Sigma, MO) to minimize clumping of the cells. For the overexpression experiments, three strains - strain of interest (SOI), Reporter, and Reference were prepared the same way, mixed thoroughly in the Eppendorf tube before loading to PDMS (polydimethylsiloxane) - based mother machine microfluidic device. The latter were prepared following a previously described procedure ^57^. The concentrated mix of three strains was pipetted onto the main flow channel of the mother machine device to populate the dead-end channels of the device for one hour. Next, a 10-ml syringe (Becton Dickinson, Fisher Scientific) was prepared with fresh M9 growth medium with the carbon source, additives, BSA (0.075 μg/ml), and appropriate antibiotics when needed and mounted on NE-1000 Syringe Pump (New Era Pump Systems, NY). The tubing was connected to the device, and the flow of fresh M9 medium started and was kept at 6 μl/min during the entire experiment. The cells were left to grow in channels overnight (at least 14 hr) to ensure steady-state growth. The next morning, two 10-ml syringes were prepared with fresh growth medium (to one syringe inducer IPTG was added) and mounted on separate NE-1000 Syringe Pumps. A T-junction was formed with three pieces of tubing and a T-connector. Before applying IPTG to induce the expression of the gene of interest (*ftsZ, ftsN, ftsA* or *ftsA*)* and Reporter (*mNeonGreen, Ypet* or *Gfp*) from the extra copy at the *attB* site (or from the plasmid), cells were imaged in regular M9 media. After 10 hours of imaging, the first pump with the regular media was turned off, and the second syringe pump containing regular media with IPTG was turned on without interrupting imaging using a custom-made LabVIEW program. Altogether, cells were imaged for 22-26 hrs. For a list of all SOI, Reference, and Reporter strains and inducer IPTG concentrations used in experiments see SI Table S4.

### Fluorescence microscopy

A Nikon Ti-E inverted fluorescence microscope (Nikon Instruments, Japan) with a 100X NA 1.40 oil immersion phase-contrast objective (Nikon Instruments, Japan) was used to image the bacteria. Images were captured on an iXon DU897 EMCCD camera (Andor Technology, Ireland) and recorded using NIS-Elements software (Nikon Instruments, Japan). Fluorophores were excited by a 200W Hg lamp through ND4 and ND8 neutral density filters. Chroma 41001 and 41004 filter cubes (Chroma Technology Corp., VT) were used to record mNeonGreen/Ypet and mCherry images, respectively. A motorized stage (Prior Scientific Inc., MA) and a Nikon Perfect Focus® system were utilized throughout time-lapse imaging. Images in M9 glucose-cas were obtained at 3 min frame rate. The typical exposure times were 400 ms at an EM gain of 200 for mNeonGreen/Ypet and 400 ms without EM gain for the phase images. The mCherry signal was recorded before the FtsN upregulation experiment to distinguish between strains JM235, STK13, and JM222 (expose time 250 ms at EM gain 200). mCherry signal was not recorded during the FtsN upregulation measurements.

### Image analysis

MATLAB, along with the Image Analysis Toolbox and DipImage Toolbox (http://www.diplib.org/) and Python 3.9 with torch and torchvision packages (https://pytorch.org/) were used for image analysis. In all analyses of time-lapse recordings, corrections to subpixel shifts between different frames were applied first. These shifts were determined by correlating phase-contrast images in adjacent frames. Individual channel images were then cropped. A convolution neural network, referred to as the Omnipose (57), was used for cell segmentation. The network was first trained on images obtained from the experimental setup used in this work. The resulting cell masks were then assembled into cell lineages using a custom Matlab script. The same script also calculated the cell lengths based on these cell masks.

### Determining induced protein concentrations

To determine the induced protein concentrations, total fluorescence intensity from fluorescent fusion proteins of the Reporter and the Reference strains were determined as a function of time from cell images. The same fluorescent fusion (either mNeonGreen or Ypet) was used in both strains. To calculate the total intensity, the pixel values from 11 pixels (1.2 μm) wide band around the cell centerline were summed. This band is slightly wider than the cell (about 0.8 μm). The intensity was then normalized per pixel value. The same normalized intensity was also calculated for the unlabeled SOI, and the average value from the cell population from SOI was subtracted from the normalized intensities from the Reporter and the Reference strain populations at each binned timepoint. The per pixel normalized and background-corrected intensity from the Reporter strain was then divided by the per pixel normalized and background-corrected intensity from the Reference strain. Since the fluorescence from the Reference did not significantly vary in time, a single global value of the per pixel normalized and background-corrected intensity from the Reference strain was used for division. The resulting quantity is referred to as the Normalized Reporter Signal and is plotted in Fig. 1D, Fig. 2A, and Fig. 3A, C in the main Text. However, this quantity reflects the induced concentration of the mature fluorescent fusion protein in the Reporter strain, not the concentration of the total fluorescent fusion proteins in the Reporter. To determine this total concentration, we fitted the data in these plots to a previously described model ^7^. The model has an analytic solution for the normalized mature protein concentration in the Reporter as a function of time:

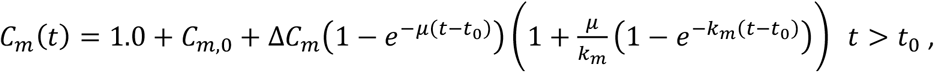

where *t*_*0*_ is the time of the induction, *C*_*m*,*0*_ is the concentration before induction due to leakage, *ΔC*_*m*_is the increase of the concentration as a result of induction, *μ* is the population-average growth rate, and *k*_*m*_is the maturation rate of the fluorescent protein. The solution before induction is *C*_*m*_*(t) =* 1.0 *+ C*_*m*,*0*_. The maturation rate in these growth conditions (glucose-CAS medium in 28 °C) for mNeonGreen was determined to be *k*_*m*_ *=* 0.029 min^-1^ previously ^7^. For Ypet we used the same chloramphenicol treatment procedure as in ^7,58^ to find *k*_*m*_ *=* 0.049 min^-1^. We calculated the population average growth rate as *μ = ln(*2)/*T*_*d*_ using *T*_*d*_ value before induction. The expression for *C*_*m*_*(t)* was fitted to the Normalized Reporter Signal using Matlab *fminsearch* function. The fittings are plotted for different proteins of interest in Fig. 1D, Fig. 2A, and Fig. 3A, C.

The fit yielded parameters *t*_*0*_, *C*_*m*,*0*_ and *ΔC*_*m*_ (the fitting line is shown by blue lines in the main text figure). Based on these parameters we calculated the total fluorophore concentration in the Reporter cells:

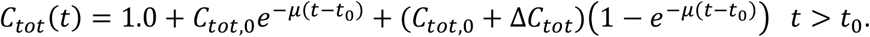

Note that *C*_*tot*,*0*_ *= C*_*m*,*0*_ and *ΔC*_*tot*_ *= ΔC*_*m*_ because both are normalized by WT concentrations. According to the above formula, the time dependence in the total fluorophore/protein of interest concentration in the cells is solely determined by the growth rate *μ*. Fitting is still needed to determine accurately *t*_*0*_, *C*_*m*,*0*_ and *ΔC*_*m*_. Since the Reporter expressed the fluorescent Reporter exactly from the same locus (the same promoter and ribosome binding site), the total fluorescent protein concentration can be taken to be equal to the total protein of interest concentration in the cells. This holds under the assumption that the protein of interest is not degraded. While FtsZ has significant degradation at lower growth rates by ClpPX protease, the degradation was undetectable in glucose-CAS medium ^7^, which is used in this work.

The excess normalized concentration of protein *X* after upregulation is defined as

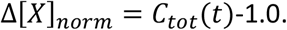

where *X* stands for FtsZ, FtsN and FtsA. *C*_*tot*_*(t) =* 1.0 corresponds to concentration in WT cells. Note that the excess normalized concentration is the excess fraction of this protein of interest relative WT level not relative to the protein level in SOI before the induction of this protein. The final excess concentration of *Δ[X]*_*norm*_is *C*_*tot*,*0*_ *+ ΔC*_*tot*_. The values for *C*_*tot*,*0*_ (and *C*_*m*,*0*_), *ΔC*_*tot*_ (and *ΔC*_*m*_) and *t*_*0*_ for each measurement are listed in Table S4.

### Determining the timing for the onset of constriction

The signal from the phase contrast images was used to determine the onset of constriction. The intensity line profiles along the long axes of the cell were averaged first, as described before ^9^. The program then searched for the local maximum in this profile near the cell middle. The constriction timing was chosen as the earliest frame when the minimum was present, with an additional condition that this minimum persisted for at least two out of three measurement frames until the cell divided.

### Modeling

We compare the experimental data to the concurrent processes model where the slowest of the two limiting processes controls the start of septum formation ^36^ rather than cell division ^44,45^. Based on an earlier report ^59^, we assume that the cell size (length) grows exponentially with rate *λ*_*bf*_ before the septum formation starts. The cells grow with a different growth rate, *λ*_*af*_, after the onset of constriction. *λ*_*bf*_ and *λ*_*af*_ is sampled from a normal distribution with the mean value determined from experimental data as 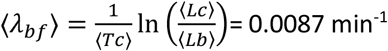 and 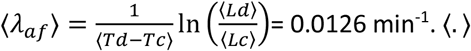 denotes the mean and the experimental data is from the 0.95x FtsZ overexpression experiment (Fig. 2A-2C). The coefficient of variation (CV) is fixed to be 0.2 consistent with the CV observed for growth rate. The two limiting processes are: 1. Inhibitory process related to DNA replication that blocks the onset of constriction for a time *Tic* that is measured from the initiation of DNA replication. *Tic* is drawn from a normal distribution with mean ⟨*Tic*⟩ and CV 0.2 which is consistent with CV values of time variables. 2. FtsZ accumulation to a threshold amount. The amount of FtsZ (number) increases at a rate proportional to the cell size i.e.,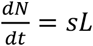. The production rate of FtsZ, *s*, is fixed at the start of each cell cycle and is assumed to be normally distributed with mean ⟨*s*⟩ *=* 1+*s*_*i*_ (in units of ⟨*λ*_*bf*_⟩) and CV 0.2. *s*_*i*_ is the leakage in FtsZ production before FtsZ is overexpressed. The upshift in FtsZ production is modeled as a step function with the mean production rate increasing from 1+*s*_*i*_ to 1+*s*_*i*_+*f* instantaneously. The factor *f* is determined using the steady-state normalized concentration of fluorescent protein in the Reporter strain. FtsZ accumulates to a threshold value drawn from a normal distribution with a mean *N*^∗^ and CV = 0.2. The values of the two model parameters ⟨*Tic*⟩ and *N*^∗^ are determined to be such that the average lengths at constriction before and after 0.95x FtsZ overexpression match between simulations and experiments. For a range of values of *N*^∗^ and ⟨*Tic*⟩, we find the value of the cost function 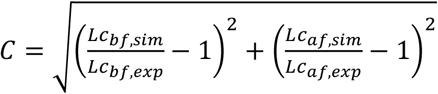 where *Lc*_*bf*,*sim*_ and *Lc*_*af*,*sim*_ are the steady-state values of *Lc* before and after overexpression obtained from simulations, respectively. *Lc*_*bf*,*exp*_ and *Lc*_*af*,*exp*_ are the steady state values of *Lc* obtained from experiments. The cost function has a minimum for ⟨*Tic*⟩ *=* 75.6 min and *N*^∗^ *=* 286 (SI Fig. S13).

To simulate the DNA replication cycle and constriction processes, we use previously specified models. The initiation of DNA replication is determined by an adder per origin model ^31,34^ where cells added a constant length increment per origin *Δ*_*ii*_ from one initiation of DNA replication to the next. *Δ*_*ii*_ is drawn from a normal distribution with mean ⟨*Δ*_*ii*_⟩ = 0.6 *μm* and CV = 0.1 determined using experimental data in previous studies ^11^. The cell division happens after a time *ΔTs* from the onset of constriction ^60^. *ΔTs* is normally distributed with a mean determined from experiments and is equal to ⟨*Td - Tc*⟩ = 18.3 min and CV is fixed to be 0.2. The value of *ΔTs* changes slightly (by 1.5 min) after FtsZ overexpression (SI Fig. S3). This is included as a step function in ⟨*ΔTs*⟩ value. The cell divides on average symmetrically but with a noise in division ratio. The standard deviation in the division ratio is 0.03 ^61^.

The upshift in FtsZ production is obtained by monitoring the concentration of a mature reporter protein. The reporter protein exists in two states-a mature state which is observed and an unmatured state. The protein goes from an unmatured to a mature state at a rate *k*_*m*_ = 0.029 min^-1^ in FtsZ upregulation measurements. The total amount of the reporter protein is a proxy for the FtsZ amount. In the simulations, the amount of mature protein (*N*_*m*_) at each time step is 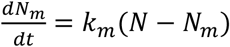, where is *N−N*_*m*_ is the amount of immature protein. Upon division, each daughter cell gets exactly half of the matured and total proteins.

We compared the results above to another model of cell division (SI Fig. S7) where the two limiting processes controlling it were: 1. FtsZ accumulation to a threshold (like in the previous model) and 2. another cell division-related protein accumulating to a threshold amount (instead of the replication-related process being limiting). The threshold amount is chosen by minimizing the cost function *C* above keeping the rest of the parameters (including FtsZ threshold *N*^∗^) fixed. We keep the noise in determining the threshold as before (CV = 0.2) and fix the threshold = 251.

## Data Availability

The datasets generated during and/or analyzed during the current study are available from the corresponding author on reasonable request.

