## Supplementary material for "Determining the rate-limiting processes for cell division in *Escherichia coli*": SI Figures and Tables

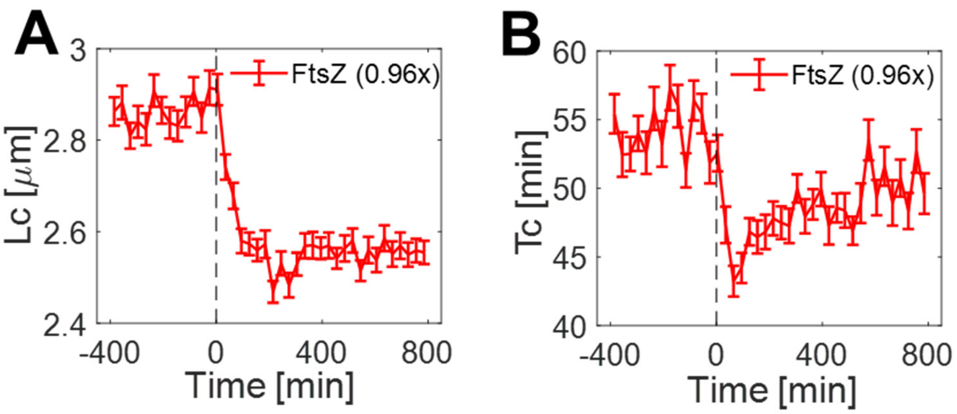


**SI Fig. S1.** Cell length and timing at the onset of constriction for the measurement shown in Fig.1 (the final $\Delta[FtsZ]_{norm} = 0.96$). (A) Cell length at the onset of constriction vs time from the upregulation of FtsZ concentration. (B) Time for the onset of constriction from cell birth vs time from the upregulation of FtsZ concentration. Time zero corresponds to adding IPTG as an inducer (indicated by a dashed vertical line). Error bars in all measurements are std errors.

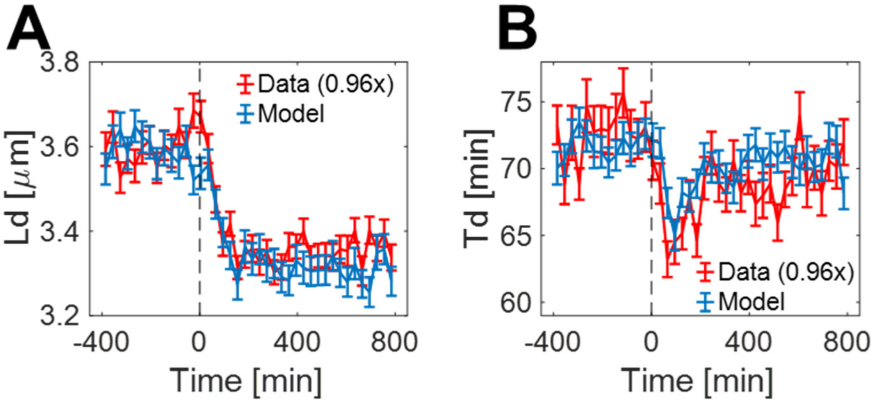


**SI Fig. S2.** Comparing experimentally measured cell length at division (A) and doubling time (B) to the predictions from the concurrent processes model (blue). The data (red) corresponds to the measurement shown in Fig.1 and Fig. 2A-C where the final $\Delta[FtsZ]_{norm} = 0.96$. The model parameters are the same as in Fig. 2A-C. All error bars are std errors.

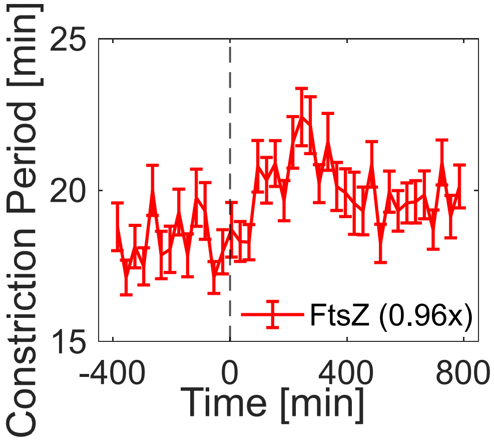


**SI Fig. S3.** The constriction period as a function of time from FtsZ upregulation (the final $\Delta[FtsZ]_{norm} = 0.96$). The constriction period is defined as $Td - Tc$. Error bars are std errors.

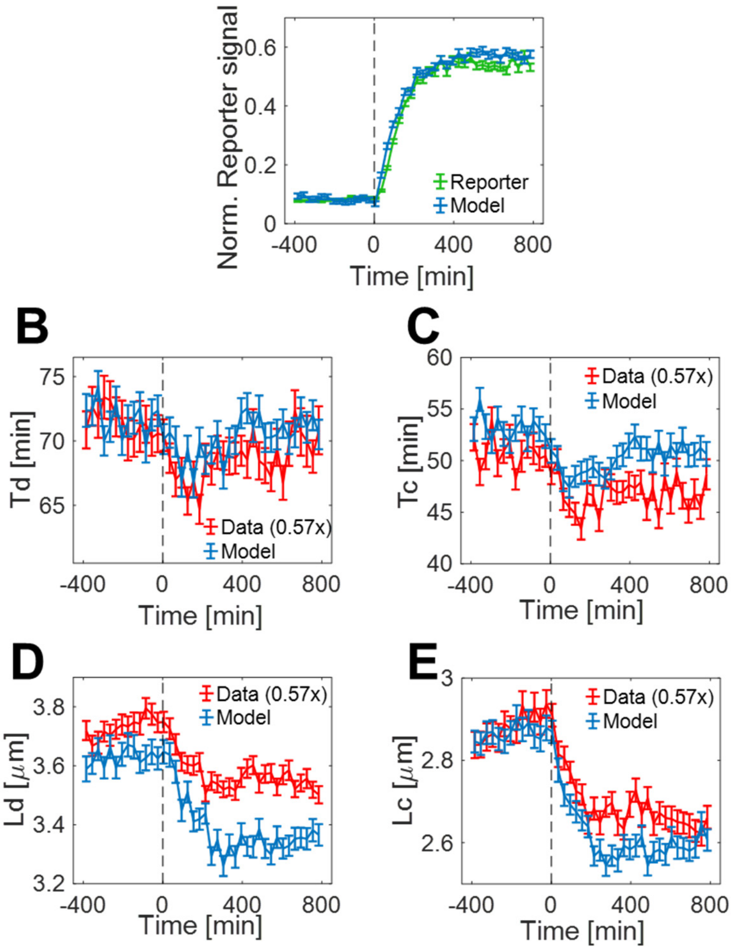


**SI Fig. S4.** Comparing experimental data (red) from the FtsZ low overexpression measurement (the final $\Delta[FtsZ]_{norm} = 0.57$) to the concurrent processes model (blue). This Figure complements the data presented in Fig. 2E of the main text. All the model parameters are the same as in Fig. 2. (A) Excess normalized FtsZ concentration vs time. (B) The doubling time vs time. (C) Timing of the onset of constriction vs time. (D) Cell length at division vs time. (E) Cell length at the onset of constriction vs time. This panel is the same as in the main text, Fig. 2E. It is reproduced here for completeness. All error bars are std errors.

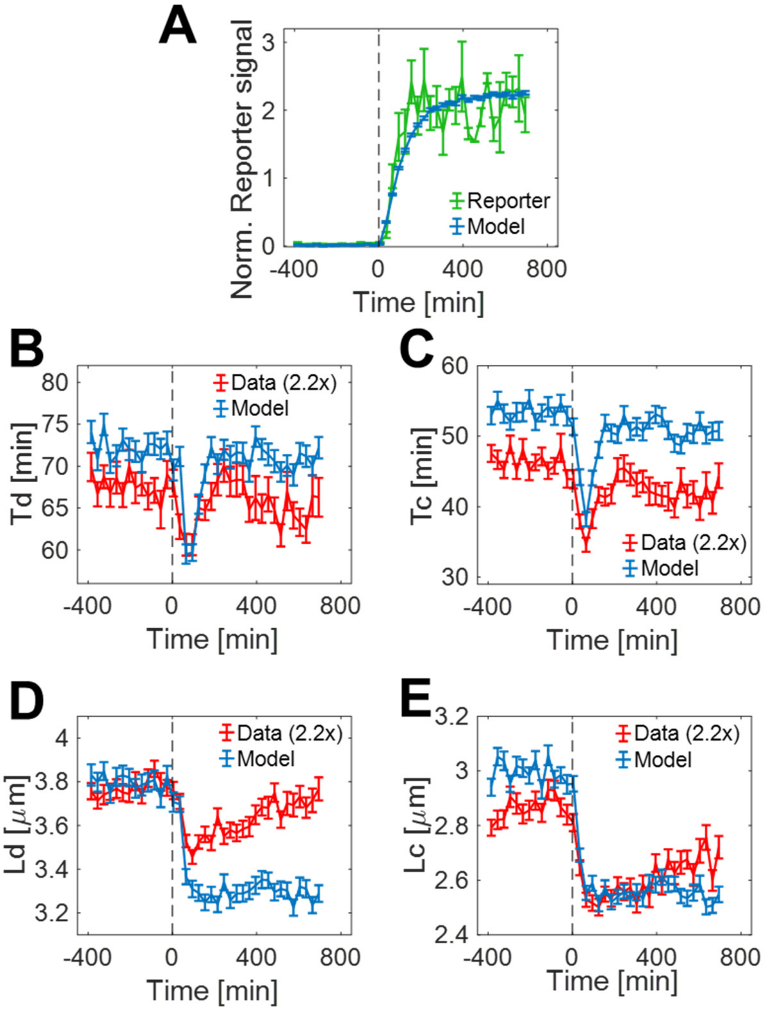


**SI Fig. S5.** Comparing experimental data (red) from the FtsZ high overexpression measurement (the final $\Delta[FtsZ]_{norm} = 2.20$) to concurrent processes model (blue). This Figure complements the data presented in Fig. 2F of the main text. All the model parameters are the same as in Fig. 2. (A) Excess normalized FtsZ concentration vs time. (B) The doubling time vs time. (C) Timing of the onset of constriction vs time. (D) Cell length at division vs time. (E) Cell length at the onset of constriction vs time. This panel is the same as in the main text Fig. 2F. It is reproduced here for completeness. All error bars are std errors.

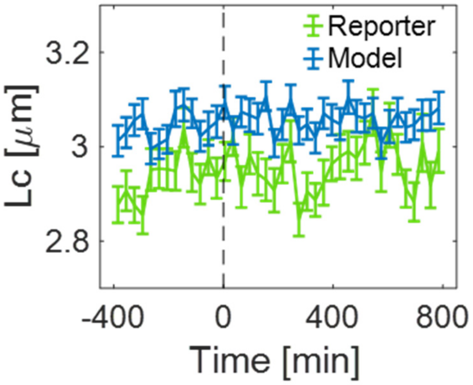


**SI Fig. S6.** Comparing experimental data on cell length at the onset of constriction from the Reporter strain (green) to the concurrent processes model (blue). The data is from the measurement with the final $\Delta[FtsZ]_{norm} = 0.96$ (shown in Fig. 1). The model parameters are the same as in Fig. 2. Error bars are std errors.

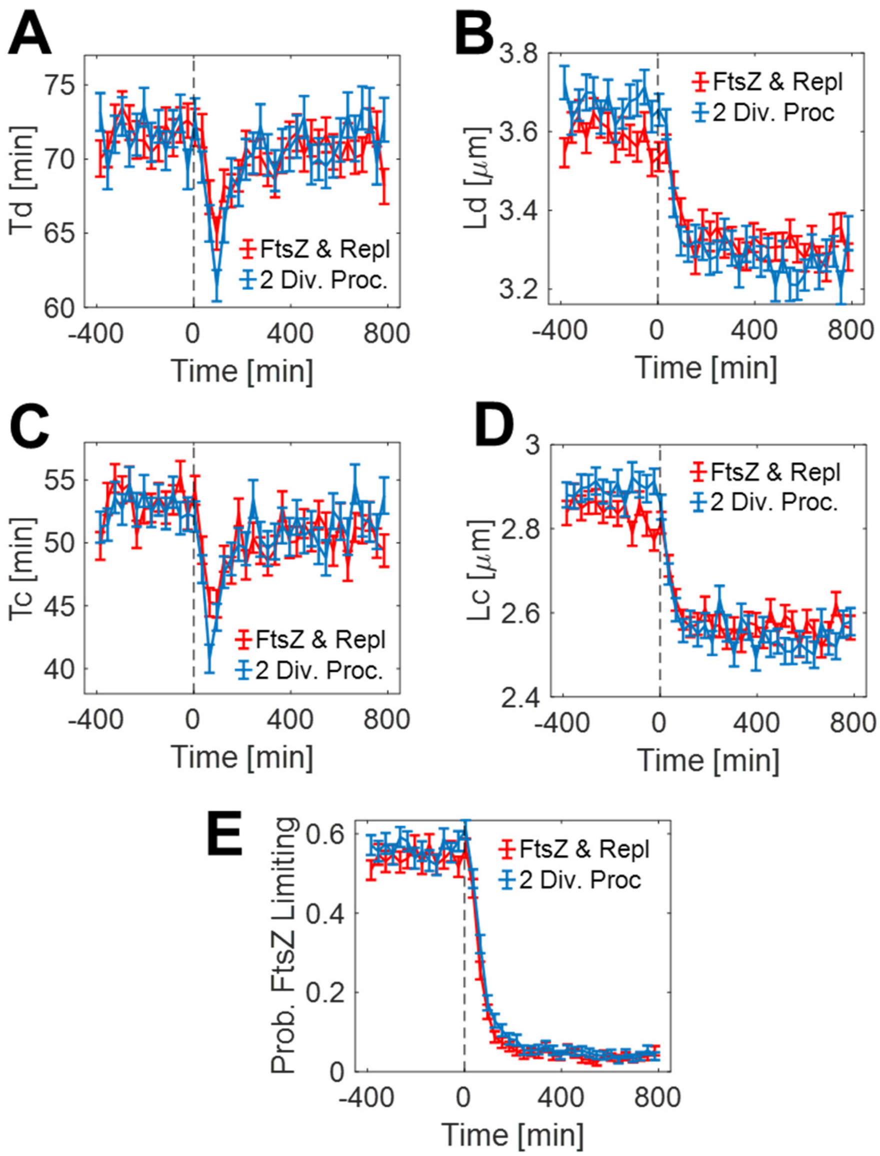


**SI Fig. S7.** Comparing concurrent processes model where FtsZ and replication are limiting (FtsZ & Repl, red) to concurrent processes model where FtsZ and some other division-related proteins (2 Div. Proc, blue) are limiting. The model corresponds to the FtsZ upregulation condition shown in Fig. 1, where the final $\Delta[FtsZ]_{norm} = 0.97$. For a detailed description of both models see Methods, *Modeling* in the main text. (A) The doubling time vs time. (B) Timing of the onset of constriction vs time. (C) Cell length at division vs time. (D) Cell length at the onset of constriction vs time. (E) Probability that FtsZ numbers in the cell are rate-limiting for the onset of constriction bases on prediction by concurrent processes model. All error bars are std errors.

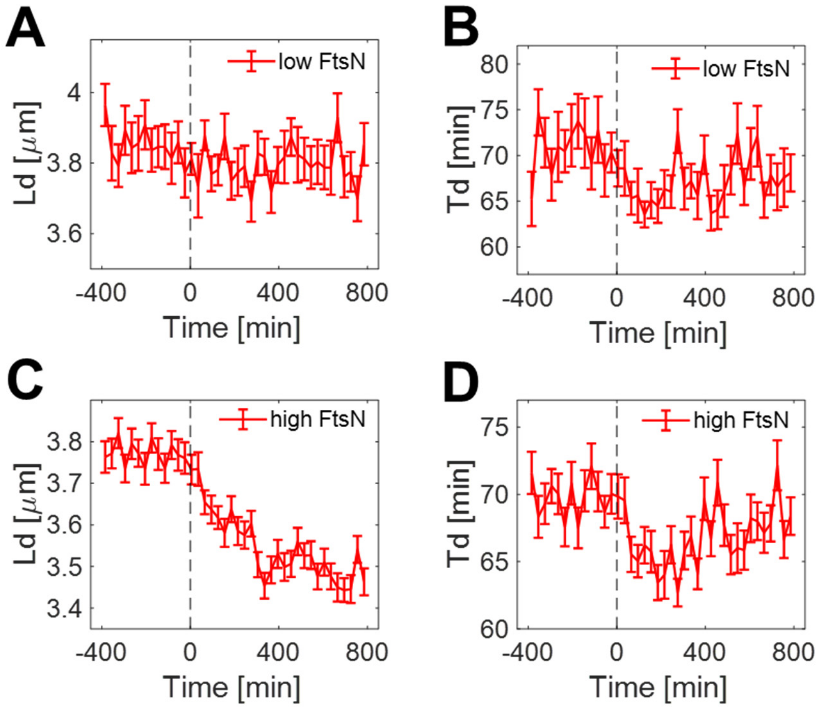


**SI Fig. S8.** Cell length at the division and the doubling time in FtsN upregulation measurements. (A-B) $Ld$ and $Td$ for the measurement where the final $\Delta[FtsN]_{norm} = 0.71$. These two panels complement Fig. 3A-C in the main text. (C-D) $Ld$ and $Td$ for the measurement where the final $\Delta[FtsN]_{norm} = 6.56$. These two panels complement Fig. 3D-F in the main text. Error bars in all measurements are std errors.

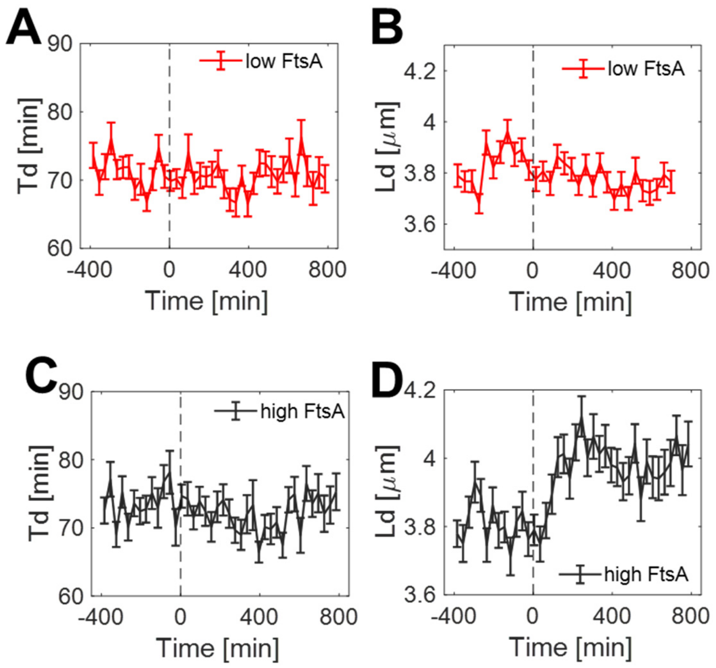


**SI Fig. S9.** Cell length at the division and the doubling time in FtsA upregulation measurements. (A-B) $Ld$ and $Td$ for the measurement where the final $\Delta[FtsA]_{norm} = 0.43$ (30 μM IPTG). These two panels complement Fig. 4B in the main text. (C-D) $Ld$ and $Td$ for the measurement where the final $\Delta[FtsA]_{norm} = 1.39$ (100 μM IPTG). These two panels complement Fig. 4C in the main text. Error bars in all measurements are std errors.

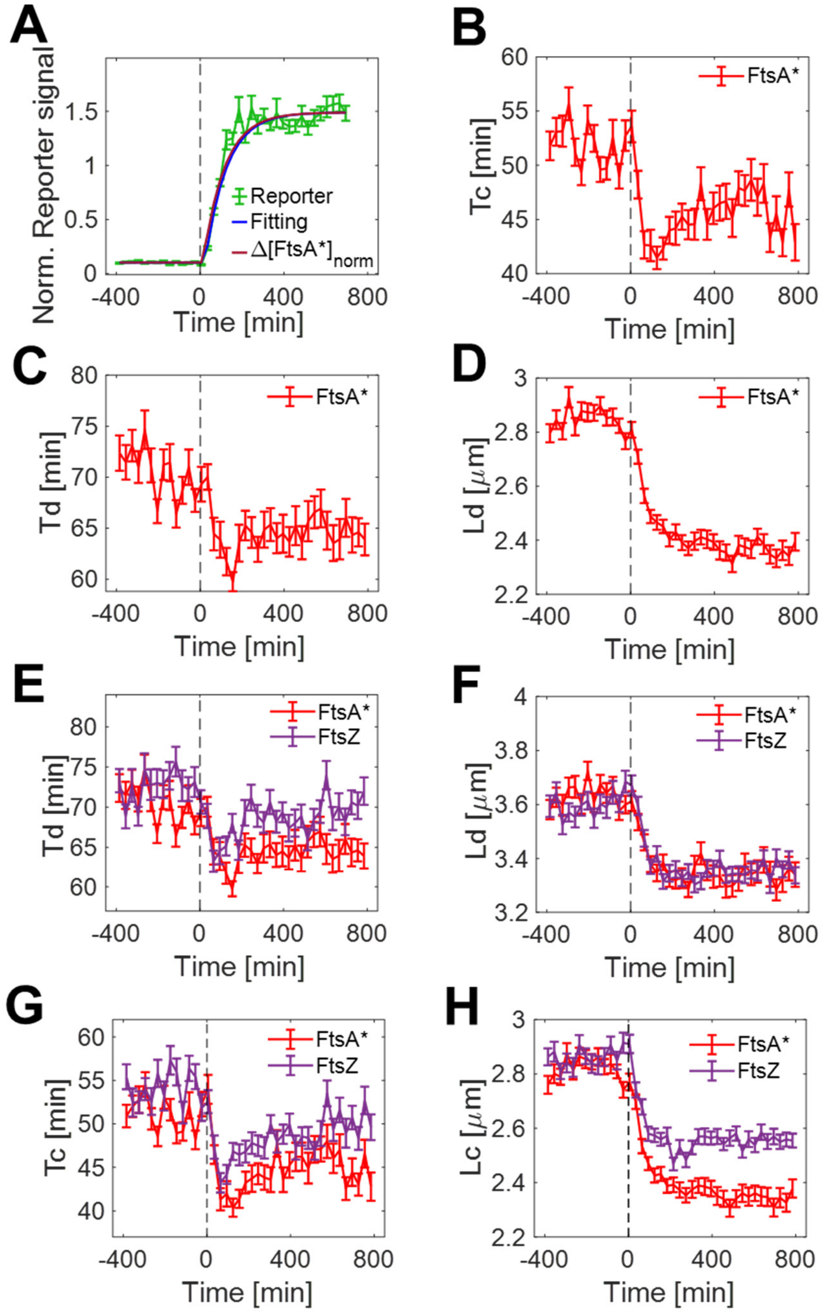


**SI Fig. S10.** Cell cycle parameters in FtsA* upregulation measurements (the final $\Delta[FtsA^{*}]_{norm} = 1.43$; 100 μM IPTG). (A) Increase of fluorescent reporter signal (Reporter) due to induction. The blue line is fitting of the model to the data and the dark red line is the calculated total upregulated protein concentration (for details, see Methods, *Determining Induced Protein Concentrations*). (B) Timing of the onset of constriction vs time. (C) The doubling time vs time. (D) The cell length at division vs time. (E) Comparing the doubling times from FtsA* upregulation measurement (red) to those from FtsZ upregulation measurement where the final $\Delta[FtsZ]_{norm} = 0.96$ (violet). (F-H) Comparing cell length at division, the timing for the onset of constriction, and cell length at the onset of constriction, respectively, from these two measurements. Error bars in all measurements are std errors.

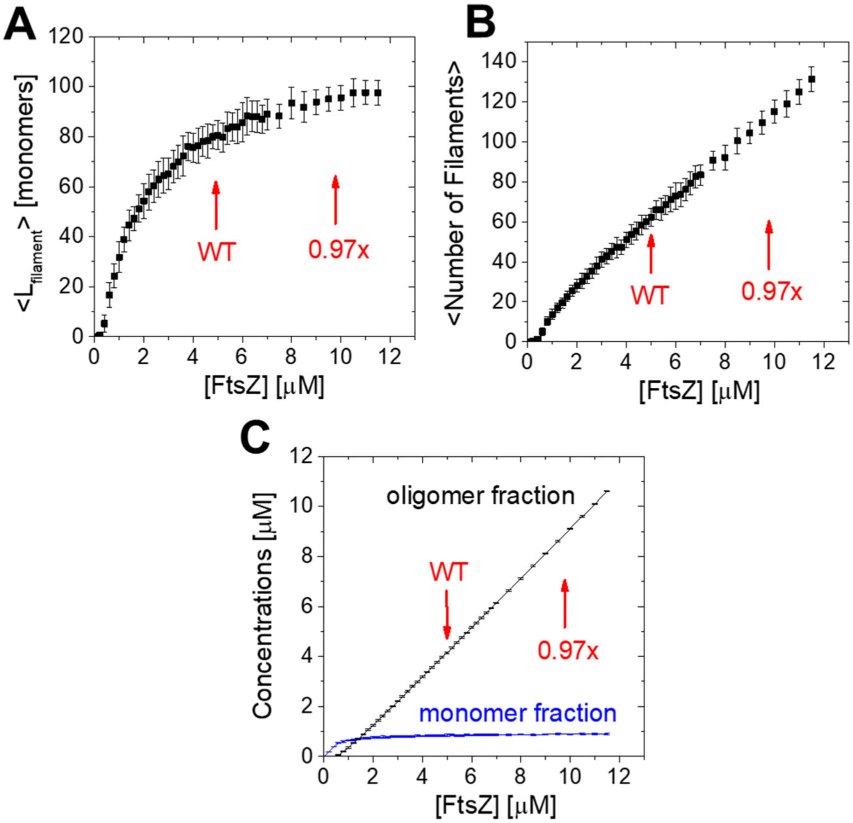


**SI Fig. S11. Modeling dependence of FtsZ protofilament length and protofilament numbers on the total concentration of FtsZ, [FtsZ], in the cell.** The used model is by Corbin and Erickson (1) and follows their parameter set for WT *E. coli*. The code and the parameter set are both available from GitHub (https://github.com/laurcor55/TreadmillModel). The cell size is taken to be 1 $\mu m^3$. The model starts from all FtsZ proteins in a monomeric form and runs for 30 seconds (real-time, not the modeling time) during which the system reaches a steady state. The values are extracted from the last time point of each modeling run and averaged over 30 runs. The error bars are std over 30 runs. The red arrow marked WT in all panels corresponds to the estimated WT concentration in the cells (5 μM), and 0.97x corresponds to the estimated FtsZ concentration in FtsZ upregulation measurements shown in Fig. 1 of the main text. (A) The average length of FtsZ protofilaments in monomer units as a function of the total FtsZ concentration in the cell. Monomers are not contributing to this average, but oligomers of length two and higher are. At the estimated WT concentration level (5 μM), the average protofilament length is 80 monomers; at 0.97x upregulated conditions, it is 96 monomers (20% increase). (B) The average protofilament number in the cell as a function of the total FtsZ concentration. At the estimated WT concentration level (5 μM), the average protofilament number is 62 monomers; at 0.97x upregulated conditions, it is 110 (77% increase). (C) The concentrations of FtsZ monomers in the monomer fraction (blue) and the polymer fraction (black) as a function of the total FtsZ concentration in the cell. The concentration of FtsZ in the polymer fraction is the monomer concentration in the polymer fraction. The critical concentration of the total FtsZ concentration where the polymer fraction appears is about 0.6 μM.

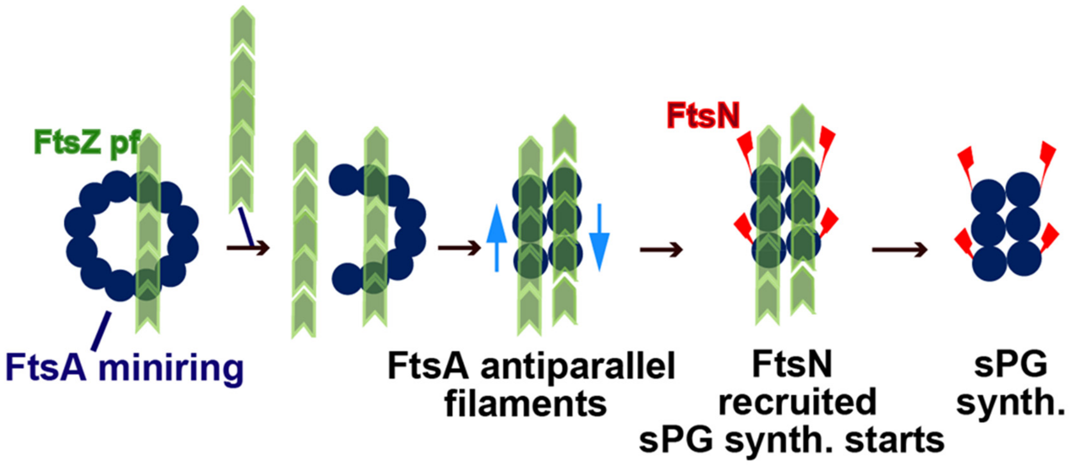


**Fig S12.** A putative mechanism where the increase in FtsZ amount in the cell leads to increased FtsZ bundling, which, in turn, can break up FtsA minirings. The broken FtsZ minirings allow the formation of FtsA antiparallel filaments, which trigger the recruitment of FtsN and the onset of constriction. The difference with models proposed earlier (2-4) is that here, the FtsZ bundling is driving these events and is not a consequence of the break-up of FtsA minirings by FtsN (3,4) or some other divisome protein (ZipA, FtsEX) (2). However, the presence of FtsA minirings is not consistent with our FtsA overexpression measurements.

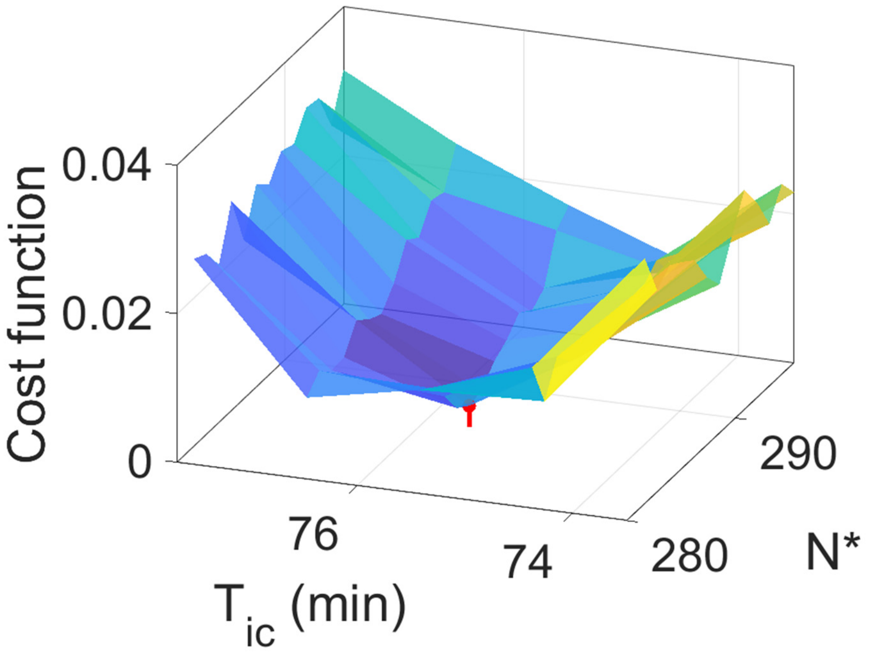


**Fig S13.** The cost function that is used in determining the model parameters $T_{ic}$ and $N^*$. $T_{ic}$ is the time from the initiation of DNA replication to the time point when the division-related process ceases to be limiting for the onset of constriction. $N^*$ is the accumulation threshold for the FtsZ molecules in the cell. The formula to calculate the cost function is given in the Modeling section of the Methods (in the main text). The minimum of the cost function occurs for $T_{ic} = 75.6$ min and $N^* = 286$ (indicated by red vertical line).

**Table S1: Strains used in the study.**

| Strain | Relevant genetic marker(s) or features | Source, reference or construction |
|---|---|---|
| **BW27783** | Δ(araD-araB)567, ΔlacZ4787(::rrnB-3), Δ(araH-araF)570(::frt), ΔaraEp-532::frt, hsdR514, ϕPcp8araE535, rph-1, Δ(rhaD-rhaB)568, λ-, | Yale Coli Genetic Stock Cente (CGSC#: 12119) |
| **BW25113** | Δ(araD-araB)567, ΔlacZ4787(::rrnB-3), hsdR514, rph-1, Δ(rhaD-rhaB)568, λ-, | Yale Coli Genetic Stock Center (CGSC#: 7636) |
| **DH5α** | F-, Φ80lacZΔM15, Δ(lacZYA-argF), U169 recA1, endA1, hsdR17(rk-, mk+), phoA, supE44, thi-1, gyrA96, relA1, λ- | Thermo Scientific competent cells |
| **HE1** | BW27783<br>ΔftsZ::ftsZ$^{55\text{-}56}$-mNeonGreen | Moore et al 2016 |
| **JM147** | BW27783<br>ΔftsZ::ftsZ$^{55\text{-}56}$-mNeonGreen<br>pEXT22 (empty vector) | Transformation of HE1 |
| **JM149** | BW27783<br>pDSW210-GFP | Mannik et al 2022 |
| **JM200** | BW27783<br>pDSW210-Ypet-FtsN | transformation |
| **JM217** | BW25113<br>pSIM5 | transformation |
| **JM221** | BW25113<br>*Δ(λattB)::lacI$^q$ P$_{Trc206}$-ypet-ftsN bla* | λ Red recombineering |
| **JM222** | BW27783<br>*Δ(λattB)::lacI$^q$ P$_{Trc206}$-ypet-ftsN bla* | P1 BW27783 X JM221, select Amp$^R$ |
| **JM228** | BW27783<br>pEXT22-mNeonGreen | transformation |
| **JM230** | BW27783<br>pDSW210-FtsN | transformation |
| **JM233** | BW25113<br>*Δ(λattB)::lacI$^q$ P$_{Trc206}$-ftsN bla* | λ Red recombineering |
| **JM235** | BW27783<br>*Δ(λattB)::lacI$^q$ P$_{Trc206}$-ftsN bla;*<br>*ΔdnaN::frt-scar-mCherry-dnaN* | P1 STK10 X JM233, select Amp$^R$ |
| **JM237** | BW27783<br>pEXT22-FtsZ | transformation |
| **JM241** | BW27783<br>pDSW210-Ypet | transformation |
| **JM242** | BW27783<br>pDSW210-FtsA | transformation |
| **JM243** | BW27783<br>pDSW210-FtsA(R286W) | transformation |

| | | |
|---|---|---|
| **JM244** | BW27783<br>*ΔftsN::frt-scar-ypet-ftsN*<br>pDSW210 (empty vector) | Transformation of STK11 |
| **JM246** | BW25113<br>*Δ(λattB)::lacI*$^{q}$ $P_{tac}$*-ftsZ aph* | λ Red recombineering |
| **JM248** | BW25113<br>*Δ(λattB)::lacI*$^{q}$ $P_{tac}$*-mNeonGreen aph* | λ Red recombineering |
| **JM250** | BW27783<br>*Δ(λattB)::lacI*$^{q}$ $P_{tac}$*-mNeonGreen aph* | P1 BW27783 X JM248, select Kan$^{R}$ |
| **JM252** | BW27783<br>*Δ(λattB)::lacI*$^{q}$ $P_{tac}$*-ftsZ aph* | P1 BW27783 X JM252, select Kan$^{R}$ |
| **STK10** | BW27783<br>*ΔdnaN::frt-scar-mCherry-dna* | Tiruvadi-Krishnan et al., 2022 |
| **STK11** | BW27783<br>*ΔftsN::frt-scar-ypet-ftsN* | Tiruvadi-Krishnan et al., 2022 |
| **STK13** | BW27783<br>*ΔftsN::frt-scar-ypet-ftsN*<br>*ΔdnaN::frt-scar-mCherry-dna* | Tiruvadi-Krishnan et al., 2022 |

**Table S2: Plasmids used in the study.**

| Plasmids | Relevant genotype | Source |
|---|---|---|
| pEXT22 | R100 ori, $P_{tac}$ promoter, $Kan^R$ | Dykxhoorn et al., 1996 |
| pEXT22-FtsZ (pJM142) | *ftsZ* in pEXT22 | This study |
| pEXT22-mNeonGreen (pJM228) | *mNeonGreen* in pEXT22 | This study |
| pDSW210 | ColE1 *ori*, $P_{206}$ promoter, $Amp^R$ | Pichoff & Lutkenhaus, 2005; Weiss *et al.* 1999 |
| pDSW210-GFP (pJM149) | *gfp* in pDSW210 | Mannik et al., 2022 |
| pDSW210-ftsA (pSEB306+) | *ftsA* in pDSW210 | Pichoff & Lutkenhaus, 2005 |
| pDSW210-ftsA(R286W) (pSEB306+*) | *ftsA(R286W)* in pDSW210 | Pichoff & Lutkenhaus, 2005 |
| pDSW210-FtsN (pJM230) | *ftsN* in pDSW210 | This study |
| pDSW210-Ypet (pJM241) | *ypet* in pDSW210 | This study |
| pSIM5 | pSC101 ori, $repA^{ts}$, $Cam^R$ | Datta et al., 2006 |

**Table S3. Oligonucleotides used in this study.**

| Name | Sequence (5’->3’) | Comment |
|---|---|---|
| PR200 | ATATTCTAGATAGAGAAAGAGGAGAAATACTAGATGTTTGAACCAATG | *ftsZ* into pEXT22 |
| PR201 | ATATAAGCTTTTAATCAGCTTGCTTACGCTG | *ftsZ* into pEXT22 |
| PR202 | ATATTCTAGATAGAGAAAGAGGAGAAATACTAGATGGTCAGCAAAGGTG | *mNeonGreen* into pEXT22 |
| PR203 | ATATAAGCTTTTACTTGTACAGTTCGTCC | *mNeonGreen* into pEXT22 |
| PR204 | ACACTGAATTCGAACTAAGGAGGATATTCATATGCC | *ypet-ftsN* (*ypet*) into pDSW210 |
| PR205 | ACGAAGCTTTCAACCCCCGGCGGCGAG | *ftsN* (*ypet-ftsN*) into pDSW210 |
| PR206 | GCTCCGGGCTATGAAATAGAAAAATGAATCCGTTGAAGCCTGCTCACTGCCCGCTTTCCA | Recombineering at λ-attachment site (*ftsZ*) |
| PR207 | GTATTAAAAACAACTTTTTGTCTTTTTACCTTCCCGTTTCGCTCTCAGAAGAACTCGTCAAGAAG | Recombineering at λ-attachment site (*ftsZ*) |
| PR208 | GCTCCGGGCTATGAAATAGAAAAATGAATCCGTTGAAGCCTGATGCATTTACGTTGACAC | Recombineering at λ-attachment site (*ftsN*) |
| PR209 | GTATTAAAAACAACTTTTTGTCTTTTTACCTTCCCGTTTCGCTCTTACCAATGCTTAATCAG | Recombineering at λ-attachment site (*ftsN*) |
| PR210 | GGCATCACGGCAATATAC | Confirmation integration at λ-attachment site |
| PR211 | GAGACACCGGCATACTCTGC | Confirmation integration at λ-attachment site |
| PR212 | TCTGGTCTGGTAGCAATG | Confirmation integration at λ-attachment site |
| PR213 | GATACCCGGGATGGCACAACGAGATTATGT | *ftsN* into pDSW210 |
| PR214 | ACGAAGCTTTTATTCGTACAATTCATTCATACCCTC | *ypet* into pDSW210 |

**Table S4. List of all measurements.** Used strains in each measurement are shown in columns SOI (Strain of Interest), Reporter, and Reference. Induction - IPTG induction level. $t_0$ –true start time of induction determined from the fitting. $t_0$ is caused by delays in fluidic lines. $C_{tot,0}$ – normalized excess concentration of protein of interest due to leakage, $\Delta C_{tot}$ – normalized excess concentration of protein of interest due to IPTG induction. The final $\Delta[X]_{norm} = C_{tot,0} + \Delta C_{tot}$ where X stands for FtsZ, FtsN, FtsA, and FtsA*.

| Measurement | SOI | Reporter | Reference | Induction [µM] | t0 [min] | C0 | dC | C0+dC |
|---|---|---|---|---|---|---|---|---|
| FtsZ Lowest | JM252<br>*N*=8643 | JM250<br>*N*=5616 | JM147<br>*N*=1758 | 30 | 13 | 0.084 | 0.49 | 0.57 |
| FtsZ Low | JM252<br>*N*=7831 | JM250<br>*N*=4956 | JM147<br>*N*=1626 | 500 | 12 | 0.097 | 0.86 | 0.96 |
| FtsZ High | JM237<br>(plasmid)<br>*N*=6956 | JM228<br>(plasmid)<br>*N*=215 | JM147<br>(plasmid)<br>*N*=44 | 100 | 12 | 0.022 | 2.18 | 2.20 |
| FtsN Low | JM235<br>*N*=2487 | JM222<br>*N*=7571 | STK13<br>*N*=3699 | 1000 | 21 | 0.068 | 0.64 | 0.71 |
| FtsN High | JM230<br>(plasmid)<br>*N*=8912 | JM241<br>(plasmid)<br>*N*=3865 | JM244<br>(plasmid)<br>*N*=2279 | 30 | 17 | 1.12 | 5.44 | 6.56 |
| FtsA Low | JM242<br>(plasmid)<br>*N*=4991 | JM149<br>(plasmid)<br>*N*=3057 | NA | 30 | 6 | 0.040 | 0.39 | 0.43 |
| FtsA High | JM242<br>(plasmid)<br>*N*=5065 | JM149<br>(plasmid)<br>*N*=1794 | NA | 100 | 7 | 0.060 | 1.33 | 1.39 |
| FtsA* | JM243<br>(plasmid)<br>*N*=6824 | JM149<br>(plasmid)<br>*N*=2961 | NA | 100 | 3.5 | 0.010 | 1.42 | 1.43 |
